# Decoding Lymphangioleiomyomatosis (LAM) Niche Environment via Integrative Analysis of Single Cell Multiomics and Spatial Transcriptomics

**DOI:** 10.1101/2025.07.07.663390

**Authors:** Ken Chen, Shuyang Zhao, Minzhe Guo, Hasan Reza, Andrew Wagner, Adnan Cihan Cakar, Cheng Jiang, Erik Zhang, Jenna Green, Emily Martin, Kathryn Wikenheiser-Brokamp, Anne Karina Perl, Debora Sinner, Jane Yu, Yan Xu

**Author notes:** These authors contributed equally to the work as first authors (Ken Chen, Shuyang Zhao). Corresponding Author Yan Xu, PhD Divisions of Pulmonary Biology and Biomedical Informatics Perinatal Institute, Cincinnati Children’s Hospital Medical Center University of Cincinnati College of Medicine 3333 Burnet Avenue, MLC 7009, Cincinnati, Ohio 45229-3039.

## Abstract

Lymphangioleiomyomatosis (LAM) is a rare, progressive lung disease characterized by cystic destruction and metastatic growth of smooth muscle-like cells. Despite advances in understanding its genetic basis, the cellular heterogeneity, regulatory mechanisms, and microenvironmental interactions driving LAM progression remain poorly defined. In this study, we employed an integrative multi-omics approach combining single-cell RNA sequencing (scRNA-seq), single-nucleus ATAC sequencing (snATAC-seq), and high-resolution spatial transcriptomics (Visium, Visium HD, and Xenium) to decode the LAM niche in its native environment. We identified two spatially and functionally distinct LAM subtypes: LAM^CORE1^ and LAM^CORE2^. LAM^CORE1^ cells exhibited a uterine smooth muscle-like phenotype, expressing associated markers (*ACTA2*, *MYH11*) and were enriched in MTORC1 signaling and myogenic pathways, supporting a uterine origin. In contrast, LAM^CORE2^ cells displayed fibroblast-like features, with upregulated extracellular matrix (ECM) remodeling genes (*COL1A1*, *MMP11*) and epithelial-to-mesenchymal transition (EMT) pathways, suggesting a role in niche formation. Pseudotime and regulon analyses revealed dynamic transitions between these subtypes, driven by distinct transcriptional networks (e.g., HOX/PBX in LAM^CORE1^, TWIST/ZEB in LAM^CORE2^). The presence of the two distinct LAM subtypes was further validated by RNAscope and immunofluorescence microscopy.

We identified LAM-associated fibroblasts (LAFs) as activated stromal cells expressing canonical markers (*FAP*, *S100A4*, *VIM, IGFBP7, SPARC*) and localized within LAM lesions. Subpopulations of LAFs, LAF-seed (proximal to LAM^CORE1^) and LAF-niche (surrounding LAM niches), exhibited unique functional profiles, including ECM deposition, TGF-β signaling, and myofibroblast activation. Regulatory network analysis pinpointed *EGR1* as a central hub governing LAF phenotype. Our comprehensive spatial profiling revealed niche structures dominated by LAM^CORE1^ cells and surrounded by lymphatic endothelial cells (LECs), LAFs, scattered LAM^CORE2^ cells, macrophages, and reprogrammed alveolar epithelial cells (AT2). ECM remodeling and aberrant organization of cable-like structures (α-smooth muscle actin+) of the LAM niches were further validated by second harmonic generation microscopy.

These findings provide a high-resolution blueprint of LAM pathogenesis, highlighting the interplay between uterine-derived LAM^CORE^ cells, activated fibroblasts, and the remodeled lung microenvironment. They significantly enhance our understanding of the LAM niche microenvironment and offer insights into potential therapeutic targets and strategies for managing this complex disease.

## Introduction

Lymphangioleiomyomatosis (LAM) is a rare metastasizing lung disease primarily affecting women of childbearing age and causes cystic remodeling of the lung and progressive respiratory failure [1, 2]. LAM is caused by deleterious mutations in *TSC1* or *TSC2*, which encode components of a hetero-oligomer that controls the activity of the mammalian target of rapamycin (mTOR) [3, 4]. The mTOR inhibitor, sirolimus, is an FDA-approved suppressive therapy that stabilizes lung function in most LAM patients [5, 6]. However, the drug does not eliminate LAM cells and can be associated with significant side-effects [5]. There is therefore an urgent need to identify new molecular targets and develop new treatments for LAM.

Uncertainty regarding the origins of LAM cells has been an obstacle to the development of new therapeutic strategies [7]. A uterine source of LAM is supported by the disease’s predominance in women, histopathological characteristics of LAM lesions consistent with uterine smooth muscle-like cells, a gradient of axial lymph node involvement from pelvis to thorax [8-10], and variation in disease progression across the female reproductive cycle [11]. Our previous single-cell study of LAM lung and uterus identified a unique population of LAM^CORE^ cells that are readily distinguishable from endogenous lung cell types and share strong transcriptomic similarity with normal uterine myocytes and uterine LAM^CORE^ cells [9]. Gene regulatory network analysis revealed activation of uterine-specific HOX-PBX transcriptional programs in pulmonary LAM^CORE^ cells [12], providing further evidence of a uterine origin. In the present study, we further identified two spatially resolved pulmonary LAM^CORE^ subtypes and characterized the dynamic evolution of uterine LAM^CORE^ cells to pulmonary LAM^CORE^ subtypes.

The tumor microenvironment is a complex and dynamic niche, comprising fibroblasts, extracellular matrix (ECM), blood vessels, immune cells, and other supporting cells. This environment plays crucial roles in promoting cancer cell survival, growth, and evasion from the immune system [13]. Cancer-associated fibroblasts (CAFs), a key component of tumor microenvironments, are instrumental in cancer progression, remodeling the ECM and secreting various cytokines, thereby recruiting other cells into the tumor stroma [14]. CAFs are defined based on a combination of morphological features, biomarkers, and genetic mutations, i.e., cells isolated from tumors that exhibit an elongated spindle morphology and express mesenchymal biomarkers such as VIM, ACTA2, IGFBP7, Fibroblast Surface Protein (FSP/S100A4), and Fibroblast Activation Protein (FAP) but lack cancer related genetic mutations [14]. CAFs are known for their significant heterogeneity and plasticity [15].

Given the similarities in the progressive, metastatic qualities of both LAM and cancer and the heterogeneous nature of the neoplasms present in both diseases, we applied cancer tumor characterization strategies to LAM lesions [16]. LAM lesions are currently characterized at the morphological level. The cellular composition, microenvironment, and cellular interactions in LAM niches remain unclear. The allele frequency of *TSC1* or *TSC2* mutations within LAM lesions ranges from 4% to 60%, with an average of less than 20% [17], supporting the hypothesis that recruited stromal cells, rather than mutation-bearing LAM cells, dominate the lesions [11, 17, 18]. LAM nodules consist of two major types of cells: spindle-shaped smooth muscle-like cells and epithelioid-like cells that are positive for PMEL (HMB45), ACTA2, and ESR antibody staining [17, 19]. LAM-associated fibroblasts (LAFs) have been identified as a population of fibroblasts within LAM nodules that express smooth muscle α-actin and CAF markers (such as S100A4, IGFBP7, CXLC12, CXCR4, and ECM) but do not express PMEL [18, 20, 21]. A recent study using proteomic profiling of late-stage LAM lung tissue showed enrichment of proteins involved in fibroblast activation and ECM deposition, suggesting that LAFs may contribute to LAM progression by facilitating ECM accumulation and remodeling of the LAM microenvironment [22]. None of the reported LAF markers are highly cell-selective, indicating the heterogeneous nature of LAFs. However, different cell states, spatial location, and dynamic regulation of LAFs, and the differences between LAFs and resident fibroblast in LAM lungs, remain largely unknown. In the present study, we employed a combination of spatial transcriptomics, single-cell RNA/ATAC sequencing, and network analysis to provide a spatially resolved comprehensive characterization of LAFs, providing a better understanding of the critical roles they play in LAM niche formation and progression.

Recent advances in spatial transcriptomic technologies enable detailed profiling of spatially resolved transcriptomes, providing a unique opportunity to interrogate cellular heterogeneity, niche microenvironment, and cellular interactions within their native environment [23, 24]. Characterizing the composition and cellular interactions of the LAM niche at single-cell resolution within its native environment has yet to be accomplished. In the present study, we integrated single-nucleus multiomics (snRNA/ATAC-seq) with spatial transcriptomics, immunohistochemistry, immunofluorescent confocal microscopy, RNAscope, and second harmonic generation microscopy to uncover cellular heterogeneity and composition within the LAM lesion microenvironment. We characterized LAM niches centered around LAM^CORE1^ and surrounded by LAM^CORE2^, LAFs, lymphatic endothelial cells, epithelial cells, capillary cells, and macrophages. Key regulators and regulatory networks controlling LAM subtypes and LAFs were predicted and cross-validated by data from different modalities and experimental approaches, revealing significant alterations in ECM remodeling, Wnt, TGF-β, and EMT signaling pathways. Our study identifies new LAM^CORE^ subtypes and provides new insights into LAM niche formation and progression, potentially leading to more effective and targeted treatments.

## Results

### Data collection and integration

For single cell transcriptomic analysis, we collected 13 single cell/single nucleus RNA-seq (sc/snRNA-seq) samples (9 previously published and 4 new) derived from female LAM patients ranging from ages 50 to 72 [12, 21, 25-27]. 16 age-matched sc/snRNA-seq female normal lung samples were extracted from the LungMAP CellRef dataset to serve as control lungs for this study [28] (**Table 1**). One previously published snRNA-seq uterine LAM sample was also used for integrative transcriptomic analysis [25].

**Table 1:** Metadata of LAM and control samples.

| PatientID | SampleID | Sex | Age | Tissue | Technology | Accession |
| --- | --- | --- | --- | --- | --- | --- |
| LAM1 | LAM1 | Female | 72 | Lung | scRNA | GSE135851:GSM4035465 |
| LAM3 | LAM2 | Female | 50 | Lung | scRNA | GSE135851:GSM4035467 |
| LAM32 | LAM3 | Female | 55 | Lung | scRNA | This study |
| LAM18 | LAM4 | Female | 69 | Lung | scRNA | This study |
| LAM_UPenn | LAM5 | Female | 70 | Lung | scRNA | GSE139819:GSM4143260 |
| LAM_UPenn | LAM6 | Female | 70 | Lung | scRNA | GSE139819:GSM4143261 |
| LAM1158 | LAM7 | Female | 52 | Lung | scRNA | GSE190260:GSM5718714 |
| LAM1163 | LAM8 | Female | 66 | Lung | scRNA | GSE190260:GSM5718715 |
| LAM1164 | LAM9 | Female | 70 | Lung | scRNA | GSE190260:GSM5718716, GSM5718717, GSM5718718 |
| LAM3 | LAM10 | Female | 50 | Lung | snRNA/snATAC | This study |
| LAM32 | LAM11 | Female | 55 | Lung | snRNA/snATAC | GSE217108:GSM6705232, GSM6705234 |
| LAM44 | LAM12 | Female | 56 | Lung | snRNA/snATAC | GSE217108:GSM6705233, GSM6705235 |
| LAM50 | LAM13 | Female | 57 | Lung | snRNA/snATAC | This study |
| LAM3 | LAM14 | Female | 50 | Lung | snATAC | This study |
| LAM4 | LAM15 | Female | 52 | Lung | snATAC | This study |
| LAM19 | LAM16 | Female | 69 | Lung | snATAC | This study |
| LAMUT | LAM17 | Female | 37 | Uterus | snRNA | GSE135851:GSM4035470 |
| LAM3 | LAM18 | Female | 50 | Lung | Visium HD | This study |
| LAM4 | LAM19 | Female | 52 | Lung | Xenium | This study |
| LAM4 | LAM20 | Female | 52 | Lung | Visium | This study |
| C51ctr | Ctrl1 | Female | 70 | Lung | snRNA | GSE171524:GSM5226574 |
| C52ctr | Ctrl2 | Female | 69 | Lung | snRNA | GSE171524:GSM5226575 |
| C54ctr | Ctrl3 | Female | 72 | Lung | snRNA | GSE171524:GSM5226577 |
| EEM-062 | Ctrl4 | Female | 57 | Lung | scRNA | LMEX0000004396 |
| EEM-067 | Ctrl5 | Female | 68 | Lung | scRNA | LMEX0000004396 |
| EEM-073 | Ctrl6 | Female | 59 | Lung | scRNA | LMEX0000004396 |
| EEM-R20 | Ctrl7 | Female | 51 | Lung | scRNA | LMEX0000004396 |
| EEM-R20 | Ctrl8 | Female | 51 | Lung | scRNA | LMEX0000004396 |
| EEM-R39 | Ctrl9 | Female | 70 | Lung | scRNA | LMEX0000004396 |
| EEM-R39 | Ctrl10 | Female | 70 | Lung | scRNA | LMEX0000004396 |
| GSM3489182 | Ctrl11 | Female | 63 | Lung | scRNA | GSE122960:GSM3489182 |
| GSM3489189 | Ctrl12 | Female | 57 | Lung | scRNA | GSE122960:GSM3489189 |
| P3_5 | Ctrl13 | Female | 51 | Lung | scRNA | EGAS00001004344, syn21041850 |
| P3_6 | Ctrl14 | Female | 51 | Lung | scRNA | EGAS00001004344, syn21041850 |
| P3_7 | Ctrl15 | Female | 51 | Lung | scRNA | EGAS00001004344, syn21041850 |
| P3_8 | Ctrl16 | Female | 51 | Lung | scRNA | EGAS00001004344, syn21041850 |

This study includes sc/snRNA-seq data from 13 LAM lung samples, 1 LAM uterus sample, and 16 control lung samples. Additionally, snATAC-seq data were obtained from 7 LAM lung samples. Spatial transcriptomic data were collected from 2 LAM lung samples, comprising 3 sequencing runs.

Seurat v4.2 RPCA was used for quality control, data integration, and batch correction. Following integration of the LAM samples, visual inspection of the integrated UMAP as well as quantitative measures to ensure that all samples, cell clusters, and technical platforms were well-represented in each cell type identified (**Supplementary** Fig. 1). The LAM samples (n=13) and control samples (n=16) were further merged and integrated to assess the alteration of genes and pathways in LAM (**Fig. 1A, Supplementary** Fig. 2).

**Figure 1:**
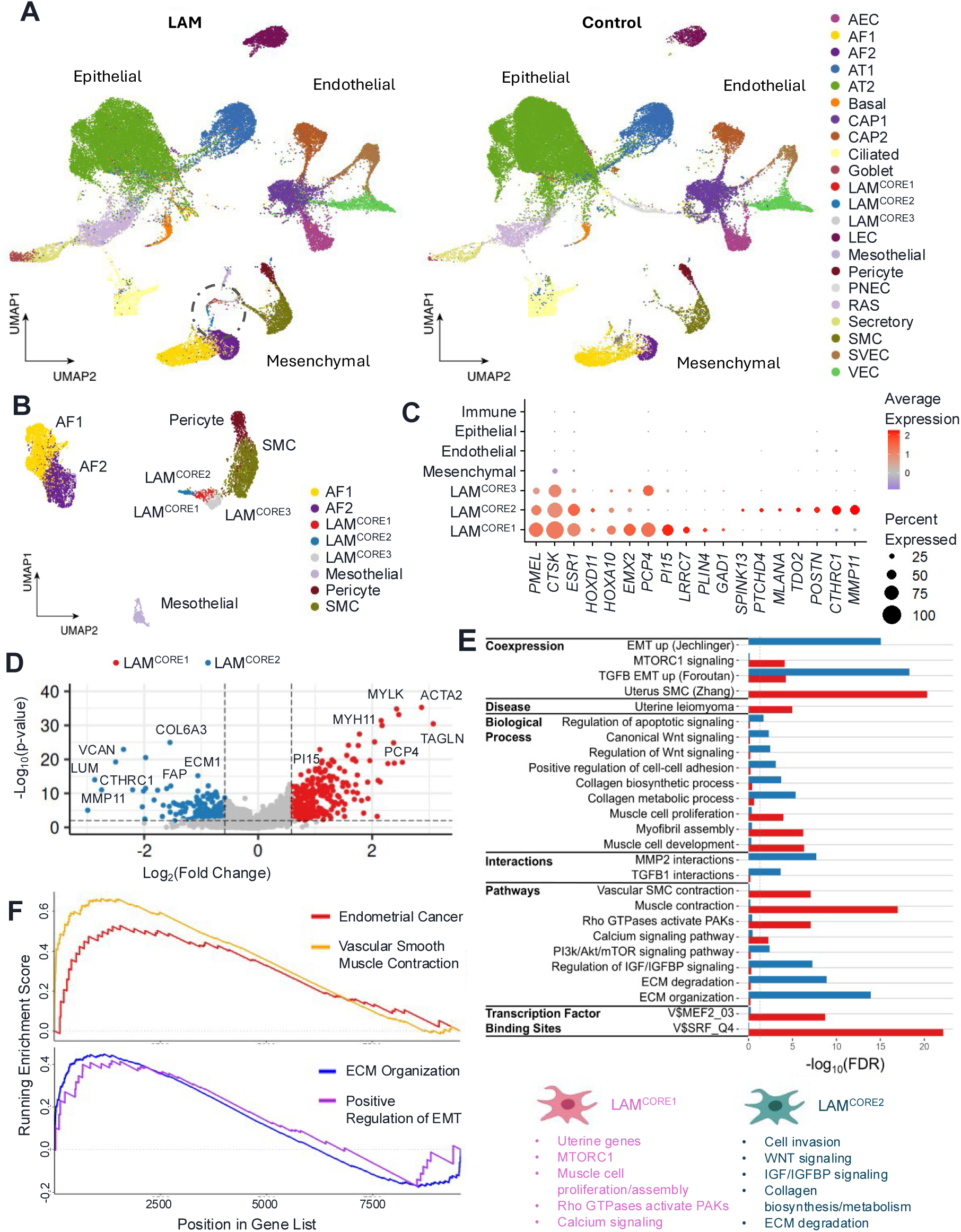
Identification and molecular characterization of LAM^CORE^ subtypes in LAM lung tissue. (A) Uniform manifold approximation and projection (UMAP) of non-immune cells from LAM and control samples, with LAM^CORE^ cells circled, highlighting their distinct clustering specifically within the LAM mesenchymal population. (B) UMAP of LAM lung mesenchymal cells used for unsupervised clustering and identification of LAM^CORE^ subtypes. (C) Dot plot of both pan-LAM^CORE^ and subtype-specific markers for LAM^CORE1^ and LAM^CORE2^, illustrating both shared and subtype-specific molecular features. (D) Volcano plot of differentially expressed genes (DEGs) between LAM^CORE1^ and LAM^CORE2^. Labeled genes illustrate distinct subtype identities. (E) Functional enrichment of LAM^CORE1^ vs. LAM^CORE2^ DEGs highlights distinct characteristics of LAM^CORE^ subtypes. (F) Gene set enrichment (GSEA) plot of select functional enrichment terms in LAM^CORE1^ (top) and LAM^CORE2^ (bottom).

### Integrative single cell data analysis identified LAM^CORE^ subtypes in the LAM lung

Our previous study identified a unique population of LAM^CORE^ cells in the LAM lung which were readily distinguished from any endogenous lung cells but shared closest transcriptomic similarity to uterine myocytes and stromal cells [25]. In addition, LAM^CORE^ cells in the lung and uterus were morphologically indistinguishable, shared similar gene expression profiles, and harbored biallelic *TSC2* mutations, supporting a potential uterine origin for LAM^CORE^ cells [25].

To further understand and assess the heterogeneity of LAM^CORE^ cells and the relationship between LAM^CORE^ cells in the LAM lung and uterus, we collected and integrated additional sc/snRNA-seq datasets from LAM and control lungs in the present study. Cell types were identified using iterative unbiased clustering analysis and automated cell type annotation using LAM Cell Atlas [27] and LungMAP Human Lung CellRef as references previously created by our group [28]. Manual curation of the cell identities was performed using selective expression of cell type markers and co-expression of signature genes at the cluster level (**Supplementary** Fig. 3-4).

In total, 39 distinct cell types were identified in the LAM lung samples (**Fig. 1A, Supplementary** Fig. 1D). These cell types were validated using known cell-type selective markers (**Supplementary** Fig. 4). Three LAM^CORE^ subtypes (LAM^CORE1^, LAM^CORE2^, LAM^CORE3^) were identified, sharing expression of highly selective known LAM markers including *PMEL, CTSK,* and *EMX2* (**Fig. 1B, C)** and were annotated as LAM^CORE^ subtypes based on the previously identified LAM^CORE^ cells in the LAM Cell Atlas reference [25, 27]. LAM^CORE3^ shared many signature genes and functions with LAM^CORE1^ but consisted of cells with a consistently low number of genes detected and lacked its own unique signature genes (**Fig. 1B**, **Supplementary** Fig. 5-6). Given these characteristics, LAM^CORE3^ is likely closely related to LAM^CORE1^, possibly representing a quiescent or less transcriptionally active state. Due to the lack of unique or differentially expressed markers, we focused our analysis on the more readily characterized LAM^CORE1^ and LAM^CORE2^ subtypes for the remainder of the study.

Since several LAM lung samples from our data collection were pre-filtered to retain only CD45^-^ cells, we removed immune cells from all samples for consistency across the dataset. The remaining LAM cells were then integrated with the previously annotated control cells for downstream analysis (**Fig. 1A, Supplementary** Fig. 2). Notably, the LAM^CORE^ subtypes were readily detectable in all LAM samples (**Supplementary** Fig. 1E) but absent in the control lung samples (**Fig. 1A**), consistent with our previous finding that LAM^CORE^ cells do not originate from endogenous lung resident cells [25].

Genes differentially expressed in the two LAM^CORE^ subtypes were identified and subject to functional enrichment analysis. As shown in **Fig. 1C**, the two LAM^CORE^ subtypes shared multiple known LAM markers such as *PMEL1* and *CTSK*, as well as novel LAM markers such as *DDR2, DES, HTR2B, KCNC2,* and *MDK* that we identified from the previous single-cell study [25]. Signature genes selectively expressed in LAM^CORE1^ cells included *PCP4, PI15, LRRC7, PLIN4,* and *GAD1*. Signature genes selectively expressed in LAM^CORE2^ cells included *MMP11, TDO2, MLANA,* and *SPINK13* (**Fig. 1C, Supplementary** Fig. 6). The complete lists of LAM^CORE^ subtype signature genes can be found in **Supplementary Table 1**.

Gene ontology and pathway enrichment analysis of LAM^CORE^ subtype signature genes identified subtype-specific and shared biological processes and pathways (**Fig. 1E-F, Supplementary Table 2**). LAM^CORE1^ displayed upregulation in muscle-related markers (*ACTA2, MYH11, MYLK,* and *TAGLN)* and functions (muscle cell development/proliferation/assembly) (**Fig. 1D-F**). Gene set enrichment analysis (GSEA) further supported this phenotype, identifying GO terms “muscle contraction” and “endometrial cancer” as positively correlated with the LAM^CORE1^ expression profile (**Fig. 1E**). Pathway analysis also corroborated these findings, predicting the activation of calcium, rho GTPases, and hallmark MTORC1 signaling pathways, all associated with smooth muscle cell behavior (**Fig. 1E)**. Interactome analysis revealed *LAMTOR5*, *MYH11, MYL6,* and *MYO18A* as major interaction hubs engaging with many LAM^CORE1^ signature genes, potentially driving and coordinating the identified pathways. Binding motifs for muscle-related transcription factors such as *SRF* and *MEF2* were over-represented in the promoter regions of the LAM^CORE1^ signature genes (**Fig. 1E**), further reinforcing the smooth muscle-like nature of LAM^CORE1^.

Importantly, the LAM^CORE1^ signature gene set showed a highly significant overlap with the transcriptomic profile of uterine smooth muscle cells (p = 4.801E-21), suggesting a strong uterine origin or lineage relationship (**Fig. 1E**). Several genes enriched in LAM^CORE1^, including *NCAM1, ITPR1, KANK1, KCNMA1, ITGA11, RBPMS, PDLIM5, THRB,* and *MYOCD*, have also been previously associated with uterine leiomyoma in genome-wide association studies (GWAS, HP_0000131; **Fig. 1E**). This molecular similarity provides additional support for the hypothesis that LAM^CORE^ cells originate from uterine-derived smooth muscle-like progenitors and retain a transcriptomic memory of their tissue of origin.

In contrast, LAM^CORE2^ signature genes showed significant overlap with signature genes of peribronchial/adventitial fibroblasts, enriched in genes associated with extracellular matrix (ECM) remodeling, collagen synthesis, and epithelial-to-mesenchymal transition (EMT) (**Fig. 1D-E**). For example, *CTHRC1* (collagen triple helix repeat containing 1) was predicted as a signature gene of LAM^CORE2^ subtypes, with expression 4-fold higher in LAM^CORE2^ than LAM^CORE1^. *CTHRC1* is a secreted glycoprotein that plays a significant role in lung ECM deposition and remodeling and is known to be associated with cell proliferation and invasion in various cancers [29-31]. Lineage tracing tools and studies in mouse models of pulmonary fibrosis identified alveolar fibroblasts as the source of *Cthrc1*-expressing pathologic fibroblasts [31]. Genes such as *COL1A1, COL6A3, ECM1, FAP,* and *MMP11* were also among the most enriched in LAM^CORE2^, suggesting robust matrix remodeling and proteolytic capacity (**Fig. 1D-E**).

Functional enrichment analysis further supported this phenotype, revealing that LAM^CORE2^ was enriched in biological processes associated with activated mesenchymal or stromal cells engaged in tissue remodeling, including ECM organization and degradation and collagen assembly, biosynthetic, and metabolic pathways (**Fig. 1E**). These processes are typically associated with activated mesenchymal or stromal cells engaged in tissue remodeling. Interactome network analysis identified *TGFB1, MMP2, FN1,* and *COL1A2* as key hubs, forming interaction networks with numerous LAM^CORE2^ signature genes (**Fig. 1E**). These genes play pivotal roles in ECM-remodeling, wound healing, and stromal activation, indicating that LAM^CORE2^ cells may adopt a state of activated mesenchymal remodeling. Moreover, genes involved in Wnt signaling, IGF/IGFBP signaling, and apoptotic regulation were upregulated in LAM^CORE2^, indicating a broader engagement of developmental and survival pathways often seen in tissue remodeling and pathological fibrosis (**Fig. 1E**). Gene set enrichment analysis (GSEA) further confirmed enrichment of ECM organization and EMT (**Fig. 1F**), processes commonly associated with tissue plasticity, cell migration, and remodeling. While EMT-related pathways were enriched, given that these cells were already of mesenchymal origin, this may more accurately reflect a transition toward a more activated, migratory, or ECM-remodeling state rather than a classical epithelial-to-mesenchymal transition.

Our data indicates that the role LAM^CORE2^ plays in LAM pathogenesis likely occurs after acclimation to the pulmonary microenvironment undergone by LAM^CORE1^, and that LAM^CORE2^ cells may play important roles in altering ECM composition and remodeling pulmonary LAM lesions. Together, these findings point to a possible transitional lineage within the LAM^CORE^ subtypes. The enrichment of uterine-specific and muscle-related features in LAM^CORE1^ strengthens the concept of a uterine origin, while the enrichment of matrix remodeling and mesenchymal activation in LAM^CORE2^ highlights the potential role of these cells in shaping the LAM microenvironment within the lung.

### The dynamic relationship between uterine LAM^CORE^ and pulmonary LAM^CORE^ subtypes

To further understand the relationship between pulmonary and uterine LAM^CORE^ cells, we integrated previously identified uterine LAM^CORE^ cells from a LAM uterus sample [25] with pulmonary LAM mesenchymal cells from the present study. The resulting UMAP displayed a significant amount of cluster-overlap and proximity between uterine LAM^CORE^ and pulmonary LAM^CORE1^ cells, with LAM^CORE2^ cells being more distal, implying a degree of transcriptomic similarity between uterine LAM^CORE^ and LAM^CORE1^ (**Fig. 2A**). The correlation scores calculated on highly variable genes were highest between uterine LAM^CORE^ and LAM^CORE1^ cells, lower with LAM^CORE2^, and negatively correlated with resident mesenchymal lung cell types (**Fig. 2B**). Expression of markers including uterine markers *UNC5D, RAMP1,* and *EMX2* and select LAM^CORE^ subtype-specific genes emphasized the similarity between uterine LAM^CORE^ and LAM^CORE1^ compared to LAM^CORE2^ (**Fig. 2C**).

**Figure 2:**
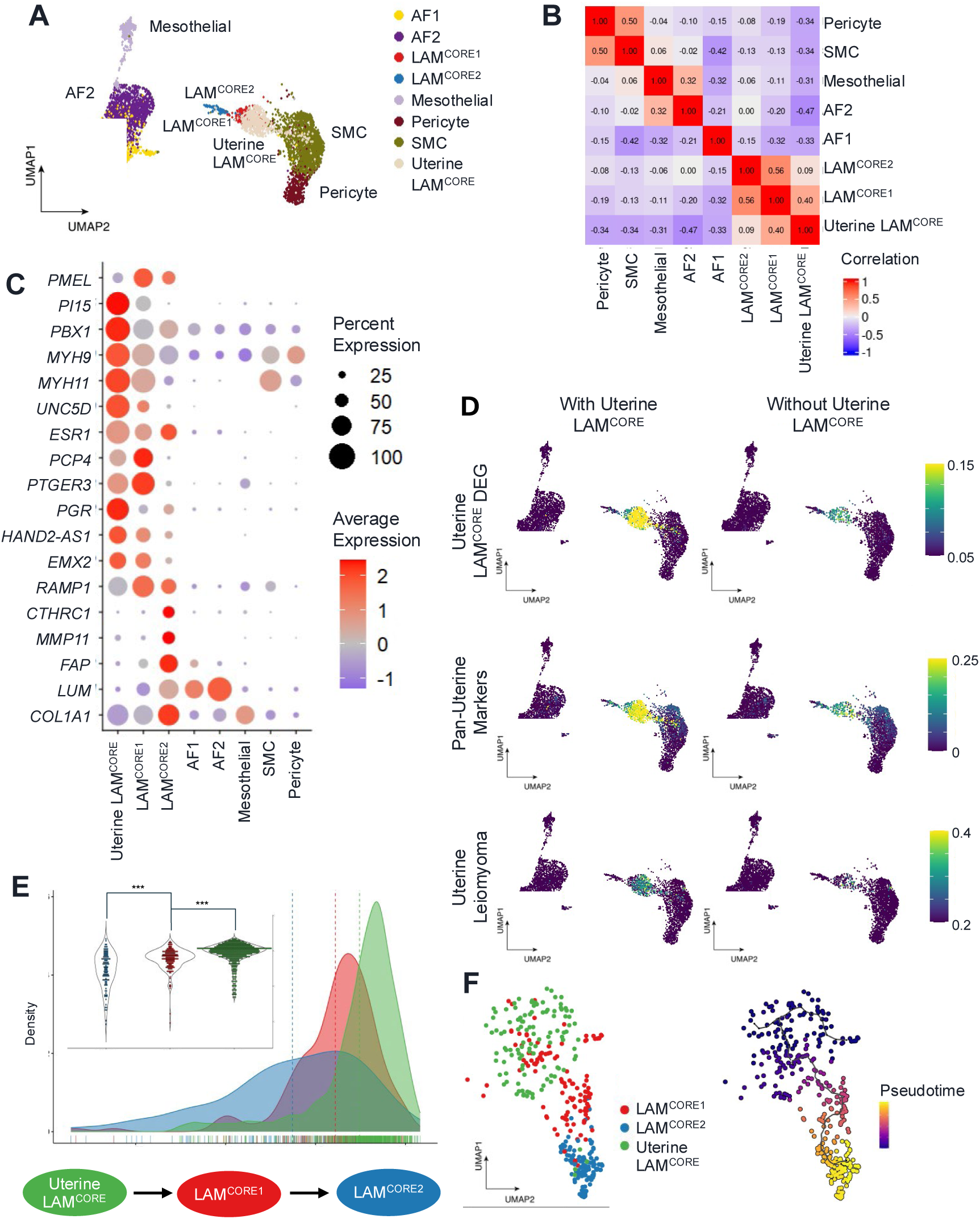
Uterine LAM^CORE^ cells resemble lung LAM^CORE1^ and highlight divergence among LAM^CORE^ subtypes. (A) UMAP of lung mesenchymal cells integrated with uterine LAM^CORE^ cells. Inclusion of uterine LAM^CORE^ emphasizes the differences between the lung LAM^CORE^ subtypes and suggests transcriptional similarity between uterine LAM^CORE^ and LAM^CORE1^. (B) Correlation heatmap of highly variable genes between lung mesenchymal cell types and uterine LAM^CORE^. (C) Dot plot showing expression of uterine-specific markers and LAM^CORE^ subtype-specific UEGs (D) UMAPs of lung mesenchymal cells with and without inclusion of uterine LAM^CORE^ cells. Uterine LAM^CORE^ DEGs were identified by comparison with other uterine cells; pan-uterine markers were obtained from GTEx bulk tissue data; uterine leiomyoma markers were derived from ToppFun GWAS data. (E) Distribution of SLICE entropy scores among LAM^CORE^ subtypes and uterine LAM^CORE^. Results suggest a transitional continuum from less mature, high-entropy uterine LAM^CORE^ to more mature, low-entropy _LAM_^CORE2^. (F) UMAP of re-clustered LAM^CORE^ cells alongside Monocle3 pseudotime trajectory analysis. Pseudotime values indicate that LAM^CORE2^ is more developmentally distant from uterine LAM^CORE^.

We previously identified the activation of a HOX/PBX centered uterine-specific transcriptional regulatory network in pulmonary LAM^CORE^ cells, which suppresses apoptosis and promotes LAM^CORE^ cell survival in the LAM lung [12]. In the present study, many of the key regulators in the HOX/PBX network (e.g., *HOXA10, HOXD10, PBX1, PBX3, RAMP1, EMX2,* and *HAND2*) were commonly induced in both LAM^CORE^ subtypes when comparing with other cell types in the LAM lungs or normal lungs [12]. Between the two LAM^CORE^ subtypes, the level of induction of these genes was generally higher in LAM^CORE1^ (**Fig. 2C)**. These patterns were also observed at a broader gene-set scale. Uterine LAM^CORE^ DEGs were identified via comparison to other resident uterine cells. These genes, along with pan-uterine markers identified through cross-tissue bulk RNA-seq comparison using data from the Genotype-Tissue Expression (GTEx) Project [32], displayed markedly higher enrichment in LAM^CORE1^ than in LAM^CORE2^ (**Fig. 2D)**. This pattern was also evident in genes found to be associated with uterine leiomyoma through GWAS (**Fig. 1E, 2D)**. This data supports the notion that the uterus is likely the source of both LAM^CORE^ subtypes with LAM^CORE1^ sharing the most proximity with uterine LAM^CORE^.

SLICE, an algorithm we developed previously to quantitatively measure cellular differentiation states based on single cell entropy, was used to assess the transitional cell states of the uterine and lung LAM^CORE^ subtypes [33]. High entropy is associated with higher functional uncertainty and more differentiation potential, while low entropy indicates more differentiated states with well-defined cell fates and functionalities. We observed statistically significant differences (p < 0.001) in mean entropy values among the three cell states, with uterine LAM^CORE^ exhibiting the highest entropy, followed by LAM^CORE1^ and then LAM^CORE2^ (**Fig. 2E**). The entropy distribution suggests a developmental track beginning with uterine LAM^CORE^ and culminating in LAM^CORE2^. Additionally, trajectory analysis assigned the lowest pseudotime scores to uterine LAM^CORE^ and the highest to LAM^CORE2^, further supporting this dynamic progression (**Fig. 2F**). The similarity between uterine LAM^CORE^ and LAM^CORE1^, along with their enrichment for muscle-related markers and functions, supports the hypothesis of a uterine origin for the disease and suggests that LAM^CORE1^ may be the result of migratory uterine LAM^CORE^ cells adapting to the pulmonary microenvironment.

Evidenced by transcriptomic proximity, uterine marker enrichment, and entropy- and pseuodotime-based analysis, a potential transitional lineage from uterine LAM^CORE^ to LAM^CORE1^ to LAM^CORE2^ becomes clear. Importantly, this stepwise transition is accompanied by a progressive shift from uterine identity and smooth muscle features to heightened ECM interaction, signaling activity, and mesenchymal remodeling. These findings suggest that LAM^CORE1^ cells may represent an initial adaptation phase, retaining uterine identity while beginning to respond to cues from the pulmonary environment and that LAM^CORE2^ cells may arise from further environmental adaptation and activation, possibly taking a more active role in creating a favorable pulmonary LAM lesion microenvironment.

### Regulon analysis identifies diverse regulatory mechanisms in pulmonary LAM^CORE^ subtypes

To further understand the regulatory circuits influencing the function and cell fate of pulmonary LAM^CORE^ subtypes, we used pySCENIC [34] and decoupler [35] to infer single-cell regulon activity. We combined the curated regulon set in the collecTRI database with those identified through pySCENIC’s co-expression and motif-based pruning algorithms and filtered to include only the 739 transcription factors (TFs) with at least five positively regulated target genes (TGs). The correlation heatmap plotted based on the regulon activity scores calculated in each cell type revealed the regulation differences between distinct cell types while maintaining intra-lineage similarity (**Fig. 3A**).

**Figure 3:**
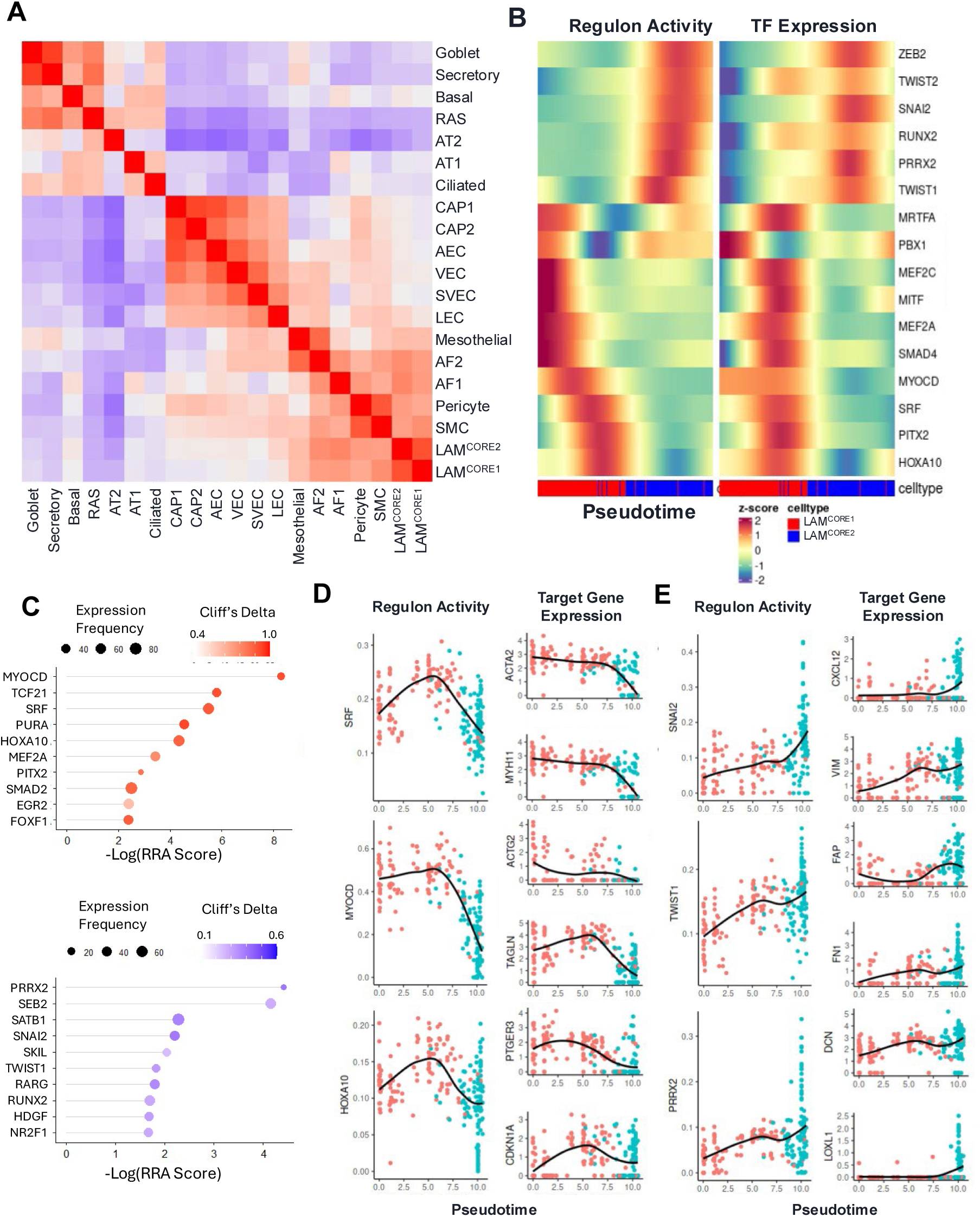
Regulon analysis reveals transcriptional drivers of LAM^CORE^ subtype identity. (A) Correlation plot of cell types using pySCENIC regulon activity scores. (B) Pseudotime plot of regulon activity and corresponding transcription factor (TF) expression in LAM^CORE1^ and LAM^CORE2^, reveals dynamic regulation across transitional trajectory and consistency between both metrics of transcriptional regulation. (C) Rankings of top regulons specific to LAM^CORE1^ (top) and LAM^CORE2^ (bottom) based on several variables, including regulon activity and transcription factor expression frequency. (D-E) Pseudotime expression profiles of select top LAM^CORE1^ (D) and LAM^CORE2^ (E) regulons and their predicted target genes, highlighting key regulators associated with each LAM^CORE^ subtype.

Differential regulon activity scores and several other variables including TF expression frequency were used to identify activated regulons that were specific to each LAM^CORE^ subtype, both in comparison to one another and to non-LAM^CORE^ cell types. These regulons were ranked using the Robust Rank Aggregation algorithm [36] to create a quantitative metric for the most enriched regulons within each LAM^CORE^ subtype (**Fig. 3C**).

Along the pseudotime trajectory, which begins with LAM^CORE1^ and transitions to LAM^CORE2^ (**Fig. 2F**), we observed a coordinated shift in regulon activity (**Fig. 3B**). Early pseudotime stages were marked by high activity of uterine and smooth muscle-related TFs including *MYOCD, SRF, HOXA10,* and *PITX2*, reflecting the uterine and myogenic nature of LAM^CORE1^. These regulons aligned with the gene expression signatures and pathway enrichments described in **Fig. 1D-F**, and their target gene sets were enriched for smooth muscle differentiation and contraction (**Fig. 3D, Supplementary** Fig. 7). Notably, genes containing SRF binding sites were previously found to be enriched in LAM^CORE1^ (**Fig. 1E**).

As pseudotime progressed toward the LAM^CORE2^ state, we observed a marked transition in regulon activity patterns, with increased activation of TFs associated with epithelial-to-mesenchymal transition (EMT), ECM remodeling, and pro-survival signaling, including *TWIST1, SNAI2, PRRX2, ZEB2,* and *RUNX2* (**Fig. 3B, E**). These regulons corresponded with a broad shift in gene expression toward ECM components (e.g., *COL1A1, COL6A3, MMP11, ECM1,* and *FAP*) and regulators of mesenchymal cell activation (**Fig. 3E, Supplementary** Fig. 8). This transition paralleled functional enrichments observed in LAM^CORE2^, including ECM organization, collagen biosynthesis, and upregulation of EMT (**Fig. 1D-F**). The increase in activity of EMT-related regulons in LAM^CORE2^ cells may reflect the emergence of a plastic, migratory mesenchymal phenotype as these cells acclimate to the lung microenvironment. Network analysis of the top LAM^CORE2^ regulons (**Supplementary** Fig. 8) revealed dense connectivity with genes modules involved in angiogenesis, ECM degradation, and apoptosis inhibition. These findings suggest that the transcriptional state of LAM^CORE2^ cells not only facilitates tissue remodeling but also promotes cellular survival and dynamic reprogramming of the surrounding stromal niche.

The pseudotime-regulon landscape reveals a temporal regulatory cascade: early activation of uterine and myogenic programs in LAM^CORE1^ is followed by a progressive shift toward EMT and ECM remodeling programs in LAM^CORE2^. This trajectory may reflect dynamic transcriptional reprogramming of LAM cells within the lung microenvironment and suggests possible regulatory checkpoints that could be leveraged for subtype-specific therapeutic targeting.

### Integrative single nucleus ATAC-seq (snATAC-seq) of LAM lungs identified LAM^CORE^ and LAM fibroblasts

The epigenetic landscape of LAM samples was examined via integrative analysis of multiome and snATAC-seq data of LAM lungs (**Table 1**) conducted with Signac [37]. snATAC-seq had less granularity in comparison with snRNA-seq but was effective at capturing major lineage and cell type distinctions. (**Fig. 4A-C, Supplementary** Fig. 9). Immune cells were removed before label transfer to maintain consistency with RNA-seq analysis. Cluster identification and annotation identified a singular LAM^CORE^ cluster, a separate endothelial and lymphatic endothelial cluster, a fibroblast cluster and several epithelial cell types (AT1, AT2, AT1/AT2, ciliated, and airway epithelial cells) (**Fig. 4A**). The LAM^CORE^ cluster exhibited selective gene activity for known LAM markers (*HOXD11, HOXD10, EMX2, HAND2*), while the fibroblast cluster showed strong gene activity of known fibroblast markers (*WNT2, TCF21*) (**Fig. 4C**). Differentially accessible peaks (DAPs) were identified and visualized, showing distinct clusters of peaks for each cell type (**Supplementary** Fig. 9).

**Figure 4:**
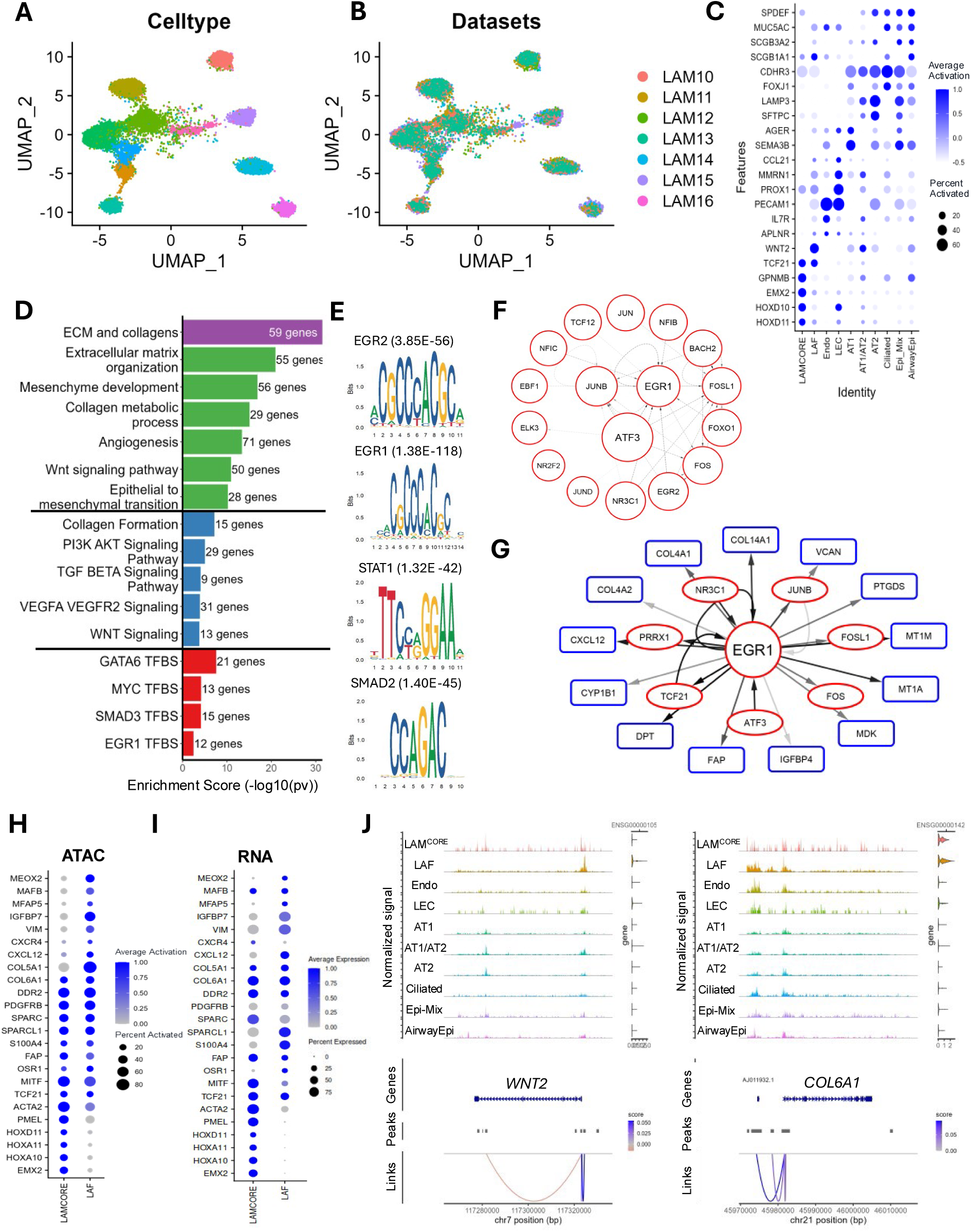
Integrative snATAC-seq and multiomic profiling uncovers characteristics of LAM lung fibroblast cells. (A-B) UMAP of integrated snATAC-seq and multiome snATAC-seq from LAM lung samples showing (A) distinct cell type clusters and (B) showing the distribution of samples across cell type clusters. (C) Dot plot of gene activity scores for canonical marker genes illustrates cell type identities inferred from chromatin accessibility. (D) Functional enrichment of genes near LAM fibroblast-specific DAPs suggests distinct pathways active in this fibroblast population. (E) Enriched transcription factor motifs (p-value < 0.05 and log2FC > 0.58) of LAM fibroblasts point to candidate regulators of cell state. (F) LAM fibroblast transcription factors (TFs) that show motif enrichment in differentially accessible peaks (DAPs) and are also identified as regulons in AF1 or AF2 subclusters via SCENIC analysis. Edges are predicted by PECA [36], with darker lines corresponding to higher confidence interactions (lower FDR). Node and text sizes are proportional to the rank of each TF from the driving force analysis, where larger nodes represent TFs with greater regulatory influence. (G) Network of selected one-hop neighbors of EGR1 from the filtered PECA network. Red circles represent TFs that are part of the broader regulatory network, while blue rectangles represent target genes that are not TFs but are predicted downstream effectors of EGR1 activity. (H) Gene activity scores for fibroblast and LAM^CORE^ markers highlight regulatory signatures within the LAM fibroblast population. (I) Gene expression from scRNA-seq data for fibroblast and LAM^CORE^ markers supports results from ATAC-inferred gene activity. (J) Peak-to-gene linkage plots for selected genes. Multiome single-cell data integrating ATAC- and RNA-seq were used to infer correlations between chromatin accessibility at distal regulatory elements and gene expression levels, identifying putative regulatory relationships.

LAM fibroblasts displayed specific upregulation of activated fibroblast markers, based on both predicted gene activity from snATAC-seq and gene expression level from sc/snRNA-seq (**Fig. 4H-I**). Importantly, these fibroblasts lacked expression of LAM^CORE^-specific markers including *PMEL, HOXD11, HOXA10* and *EMX2* (**Fig 4H-I)**. LAM fibroblasts likely include an activated LAM-associated fibroblast (LAF) population based on the characteristics of LAF described previously [18, 20, 21].

Functional enrichment analysis of LAM fibroblast DAPs revealed strong activation of canonical fibroblast programs, including ECM organization, collagen biosynthesis, and mesenchymal development, all hallmarks of activated fibroblasts (**Fig. 4D**). Notably, enrichment of cancer- and stem cell-associated pathways such as EMT, angiogenesis, and Wnt, TGF-β, and PI3K-AKT signaling suggests that these fibroblasts may adopt a pro-tumorigenic and regenerative phenotype reminiscent of cancer-associated fibroblasts (**Fig. 4D**). The presence of MYC binding site enrichment further supports a transcriptional landscape favoring proliferation and cellular plasticity. Transcription factor motif enrichment analysis identified *STAT1, SMAD2, EGR1,* and *EGR2* as key regulators of chromatin accessibility in these cells (**Fig. 4E**), several of which are known to play roles in fibroblast activation and response to TGF-β signaling [38, 39]. Together, these findings highlight the potential of LAM fibroblasts to function as activated, niche-shaping cells that contribute to disease progression through ECM remodeling and paracrine signaling.

### Gene Regulatory Network (GRN) regulating LAM Fibroblasts

A gene regulatory network (GRN) was generated by applying the PECA2 [40] algorithm to pseudobulk RNA- and ATAC-seq data from LAM fibroblasts. To prioritize biologically relevant transcription factors (TFs), we compared the GRN TFs with those identified through regulon and motif enrichment analyses (**Fig. 3**, **Fig. 4E**). TFs common to all three approaches were ranked using SINCERA driving force analysis [41], which highlighted *EGR1* as a key regulatory hub in the LAM fibroblast population (**Fig. 4F**).

The *EGR1*-centered GRN revealed direct interactions with TFs such as *ATF3*, *PRRX1*, and *NR3C1*, suggesting their potential role as co-factors in modulating LAM fibroblast-specific transcriptional programs (**Fig. 5G**). Downstream targets of *EGR1* include known markers of activated fibroblast (*FAP*, *CXCL12*, and *IGFBP4*), ECM-related genes (*COL4A1*, *COL14A1*), and pro-inflammatory mediators (*JUNB*, *FOS*, *FOSL1*), collectively pointing to *EGR1* as a driver of both structural remodeling and inflammatory signaling in the LAM niche.

**Figure 5.**
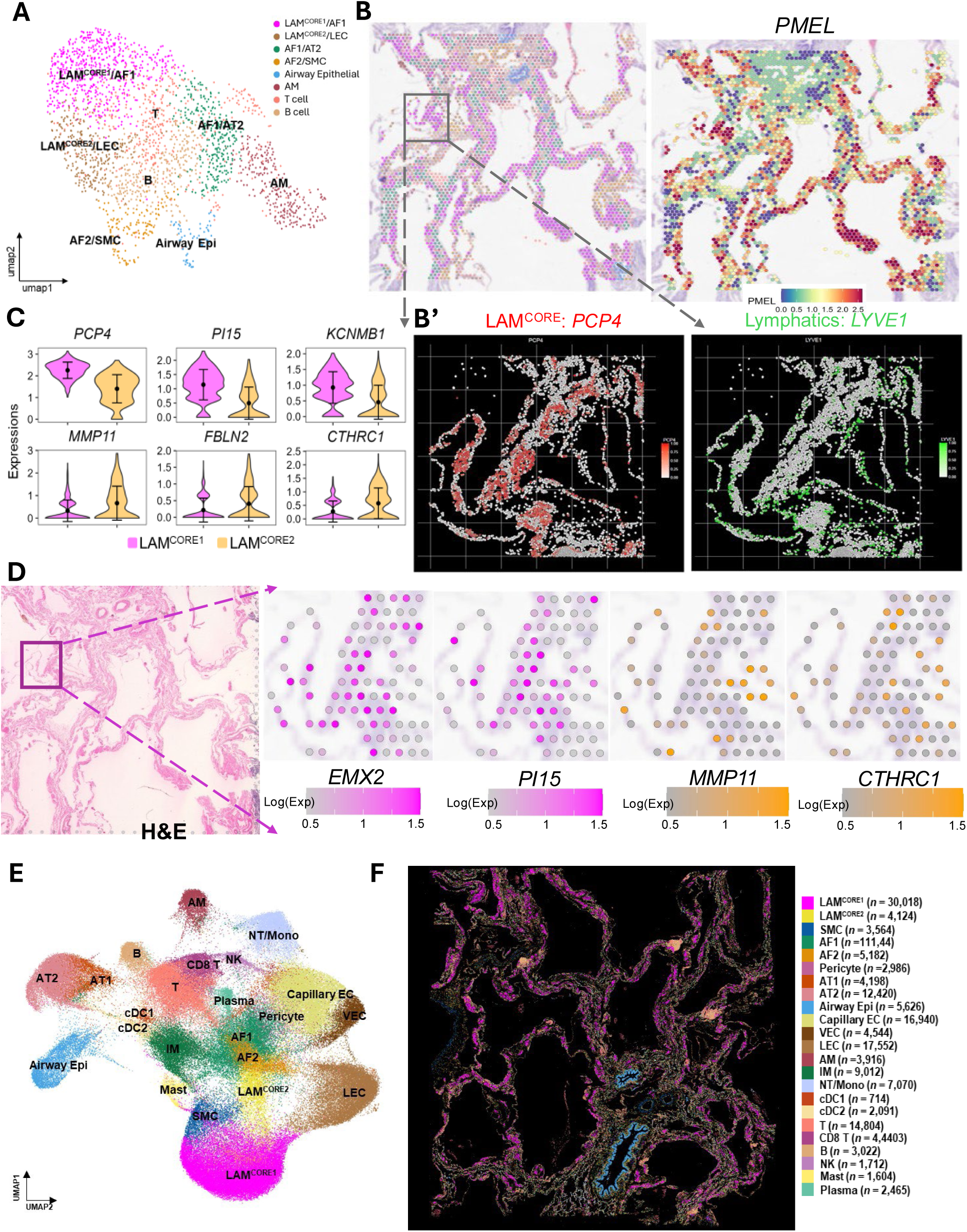
Identification and characterization of LAM^CORE^ cell subtypes via 10x Visium and Xenium. (A) UMAP visualization of identification of distinct cell populations using Visium data. (B) Spatial mapping of all identified cell types and expression of PMEL overlaid on Visium H&E. (B’) Expression of select LAM^CORE^ and lymphatic endothelial markers in the corresponding region using Xenium. (C) Violin plot showing the expression of selected LAM^CORE1^ and LAM^CORE2^ markers. (D) LAM^CORE1^ and LAM^CORE2^ markers expressions overlaid on the Visium H&E. (E) UMAP plot of Xenium spatial transcriptomic data of LAM lung. Cells were colored by cell types. (F) Spatial mapping of all identified cell types overlaid on tissue morphology, showing anatomical localization of diverse populations.

To assess the regulatory potential of chromatin-accessible regions, peak-to-gene linkage analysis identified enhancer-like regions upstream of *WNT2*, *COL6A1* and *IGFBP7*, correlating with elevated gene expression in LAM fibroblasts (**Fig. 4J, Supplementary** Fig. 10). Similar enhancer associations were observed for additional activated fibroblast markers such as *FAP*, *PDGFRB*, and *SPARC*. These findings underscore the importance of epigenetic remodeling in establishing LAM fibroblast identity and highlight enhancer activity as a mechanism contributing to the dysregulated transcriptional landscape in LAM fibroblasts.

### Validation of LAM^CORE^ subtypes via spatial transcriptomics

In our study, we utilized a combination of 10x Visium, Visium HD and Xenium to characterize LAM subtypes in their native environment (**Table 1**). The Visium platform offers whole-transcriptome coverage with a spot resolution of 55 µm, allowing for broad spatial distribution analysis. In contrast, the Xenium platform provides subcellular resolution with high capture sensitivity, focusing on a panel of 377 human cancer-related genes pre-determined by 10x Genomics. This combination enabled us to perform comprehensive analysis of the same LAM lung region, revealing both whole transcriptome spatial distribution and single-cell resolution in the LAM niches. In addition, we applied Visium HD on a different LAM patient sample for independent niche assessment and validation. Visium HD is the newly released spatial platform capturing whole transcriptome data at single-cell scale, ideal for spatial discovery and validation of single-cell multiomic predictions.

Using CellRef [28] and the LAM Cell Atlas [27] as references, we mapped cell types within the LAM lesion via robust cell type decomposition (RCTD) [42]. Visium data revealed 8 cell clusters including a large cluster of LAM^CORE1^ cells (**Fig. 5A**), characterized by high expression of *PMEL*+ spots (**Fig. 5B**). Some Visium clusters consisted of mixed cell type annotations which shared spatial proximity, such as LAM^CORE1^ /alveolar fibroblast type 1 (AF1) and LAM^CORE2^ /lymphatic endothelial (LEC), likely due to the low spot resolution of Visium. The corresponding zoomed-in region of a representative LAM niche on the Visium and Xenium platforms confirmed the presence of positive LAM^CORE^ cells (red, *PCP4*+) and abundant LECs (green, *LYVE1*+) lining lymphatic channels in the niche (**Fig. 5B’**). We gained further cell type resolution using the extended marker list from the corresponding Visium data within the region of interest. As shown in **Fig. 5C-D**, the distribution of LAM^CORE1^ and LAM^CORE2^ cells can be determined using the spatial expression of multiple LAM^CORE^ subtype markers identified by single cell analysis.

Additionally, data from Xenium identified 28 distinct cell types, with clear separation of subtypes such as LAM^CORE1^ and LAM^CORE2^, AF1 and AF2, alveolar type 1 (AT1) and AT2 (**Fig. 5E-F**). LAM^CORE1^ cells were more abundant than LAM^CORE2^ in the LAM lesion and formed dense clusters along the alveolar wall, while LAM^CORE2^ cells displayed a more scattered, single layer-like distribution (**Fig. 6 A-C**). The pairwise design utilizing both 10x Visium and Xenium platforms allowed us to leverage the strengths of each: the comprehensive whole transcriptome coverage from Visium and the high capture sensitivity and subcellular resolution of Xenium. This approach provides a detailed understanding of the cellular heterogeneity and spatial organization within the LAM lesion microenvironment.

**Figure 6.**
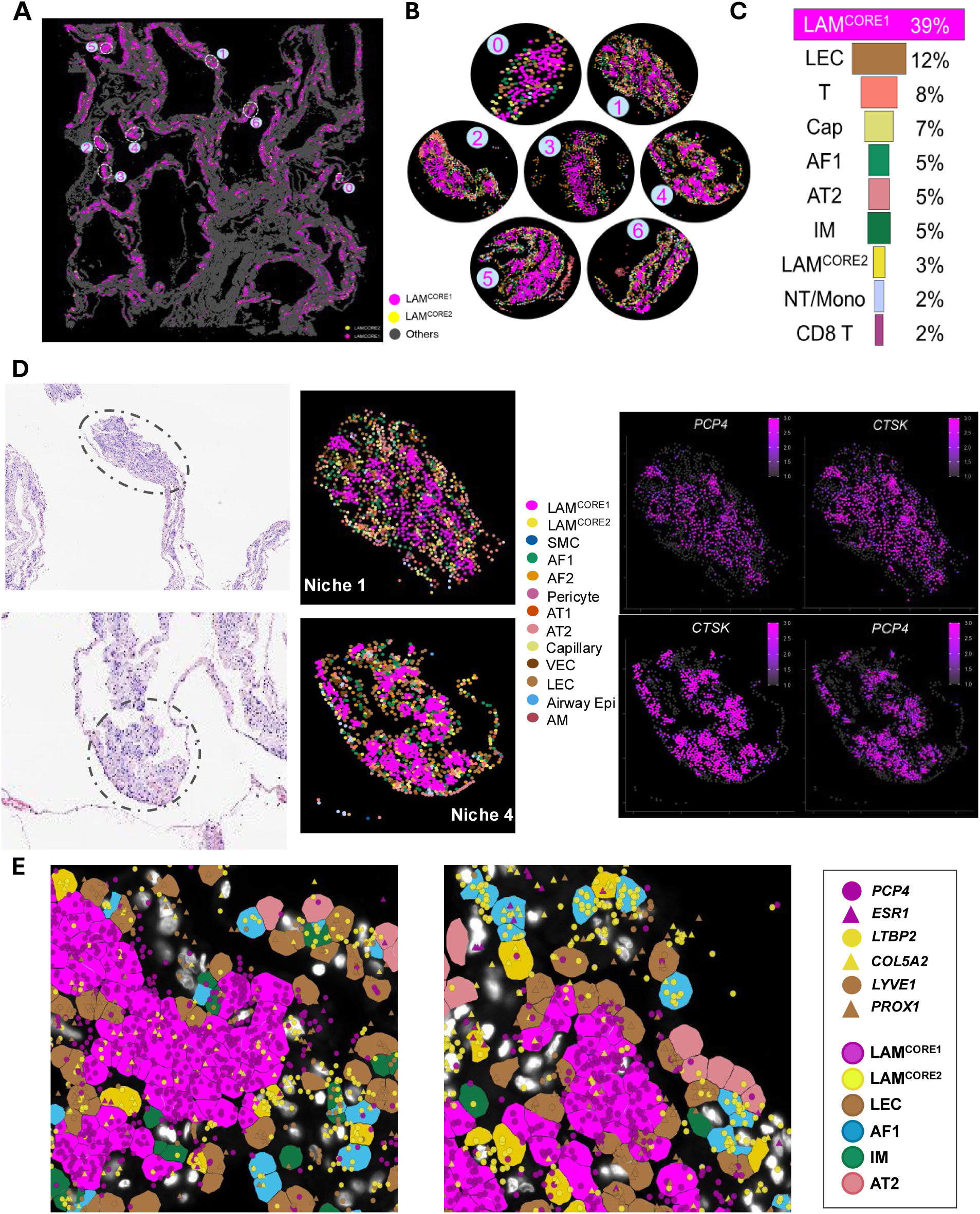
Identification and Characterization of LAM Niches using Xenium spatial profiling. (A) Spatial plot showing the automated identification of LAM^CORE^ (LAM^CORE1^ magenta, LAM^CORE2^ yellow) niches. Seven specific regions were then selected for further assessment. (B) Cell population showing across individual selected spatial niches. (C) Bar plot summarizes the average proportions of the top 10 cell types across the identified LAM^CORE^ regions, illustrating the cellular diversity and relative abundance of LAM^CORE1^ and associated populations. (D) Zoomed-in view of Niches 1 and 4, colored by cell type alongside corresponding histological images. Both niches display significant expression of the LAM^CORE^ marker CTSK and PCP4, supporting their molecular and spatial characterization. (E) Close-up view of Niche 1 cell boundaries using Xenium Explorer.

### Profiling of LAM niches using high-definition spatial transcriptomics

Considering the relatively low resolution of Visium spots, we choose to rely on Visium HD and Xenium, the technologies with single-cell subcellular resolution to identify and characterize LAM niches. Using Seurat’s BuildNicheAssay function on Xenium data, we unbiasedly identified the most LAM^CORE^ enriched clusters. To dissect the cellular composition of LAM niches, we randomly selected seven spots within LAM^CORE^-enriched clusters for detailed spatial analysis (**Fig. 6A-B**). LAM^CORE1^ cells constituted the predominant population (39%) within these niches, forming characteristic nodular or cable-like structures (**Fig. 6C**). These central clusters were surrounded by a heterogeneous microenvironment comprising recruited lung-resident cell types, including LECs (12%), T cells (8%), AF1 (5%), AT2 (5%), capillary cells (7%), and interstitial macrophages (IM; 5%) (**Fig. 6C**). The prominence of LECs, a histopathological hallmark of LAM, aligns with clinical observations of lymphatic channel proliferation adjacent to cystic lesions. Notably, LAM^CORE2^ cells were consistently present but less abundant (3% on average) and distributed at the niche periphery (**Fig. 6A-D**). High-resolution imaging of Niche 1 revealed a tightly organized core of LAM^CORE1^ cells, surrounded by LECs (*LYVE1+/PROX1+*) in direct contact with the lesion cell clusters. The niche periphery comprised AF1, AT2 cells, and macrophages, while scattered LAM^CORE2^ cells (expressing *LTBP1*, *COL5A2*) often were in direct contact with LAM^CORE1^ cells and/or nearby fibroblasts (**Fig. 6E)**. These spatial relationships support a model wherein LAM^CORE1^ cells act as uterine-derived seeds, potentially disseminating via lymphatic routes, while LAM^CORE2^ cells interact with LAFs, driving stromal activation and ECM remodeling at the niche periphery.

LAM niche and cell composition analysis was independently assessed using Visium HD on a different LAM lung sample. Spot deconvolution via RCTD was used to classify and label bins with cell types derived from the LAM Cell Atlas reference [27]. A customized script was developed for automated LAM niche detection with up to 50 µms from LAM^CORE^ cells defined as peripheral region (modified from [43], detail see Method) (**Fig. 7A-B)**. We randomly selected 10 LAM^CORE^ enriched niche spots (**Fig. 7, Supplementary** Fig. 11). Consistent with results from Xenium-based analysis, LAM^CORE^-enriched niches were organized with LAM^CORE1^ cells forming a dense core (32%) surrounded by AF1 (10%), AT2 cells (9%), LAM^CORE2^ (8%) and LECs (5%) (**Fig. 7C-D**). This concentric pattern suggests that LAM^CORE1^ cells may serve as a structural or signaling hub within the lesion. In contrast, LAM^CORE2^ cells present in each of the niches (5-12%) display more scattered and peripheral distributions, likely forming more and closer interactions with surrounding resident lung cells in the niche microenvironment (**Fig. 7F, Supplementary** Fig. 11). Considering the predominant and central locations of LAM^CORE1^ cells in the LAM niches, we measured distances from each LAM^CORE1^ cell to major cell types within the predicted niches. LAM^CORE1^ cells were in closest proximity with LECs (**Fig. 7H**). Consistent with Xenium observations, LECs exhibited more direct contact with LAM^CORE1^ cells while AF1 and AT2 cells were more peripheral (**Fig. 6E, 7H**).

**Figure 7.**
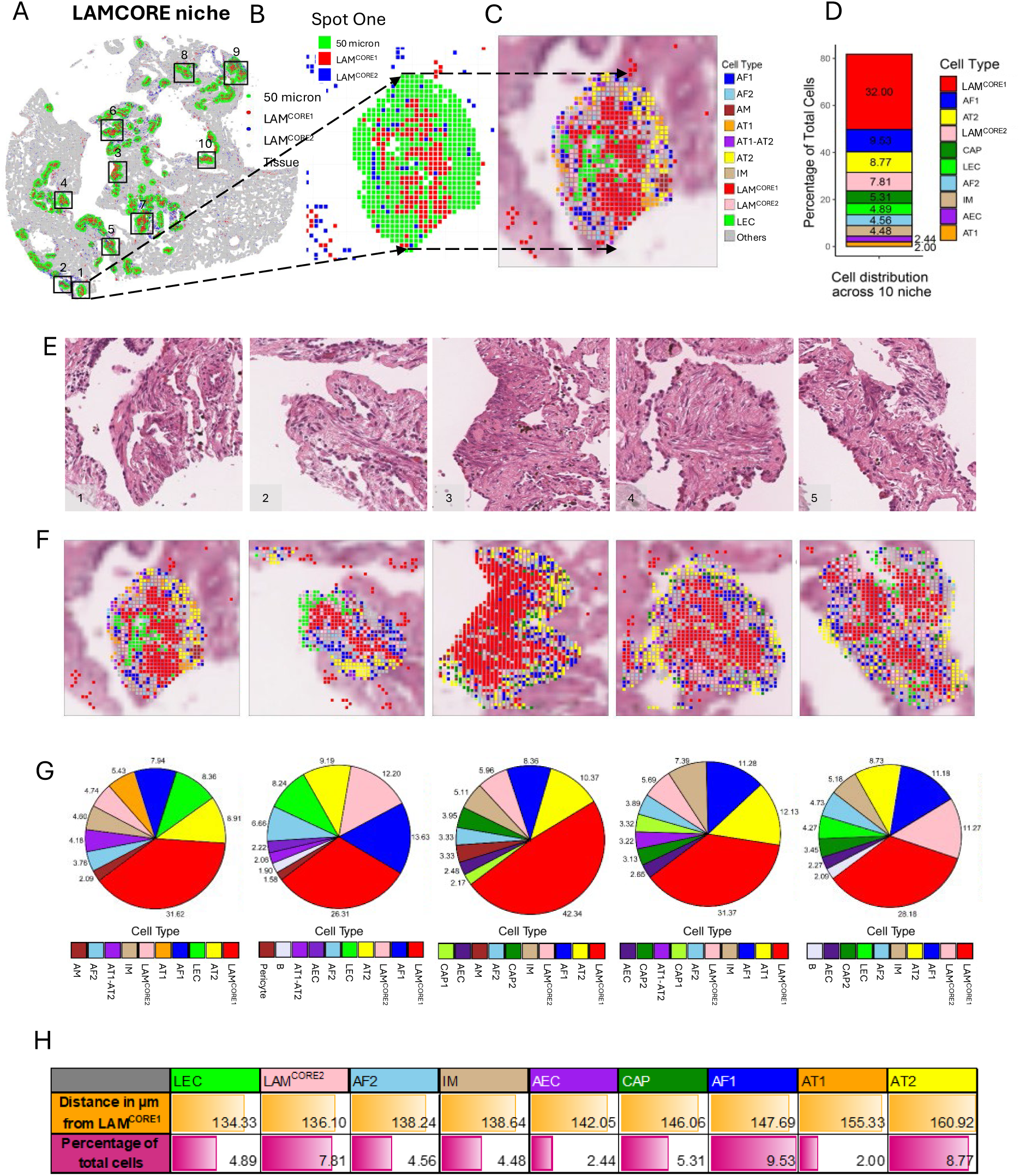
Identification and Characterization of LAM Niches using Visium HD. (A) Spatial mappings of LAM^CORE^ enriched niches. (B) Niche 1 as a representative for each LAM^CORE^ subtype. (C) Cell types identified in Niche 1. (D) Average cell type distribution across all LAM^CORE^ niches identified. (E-G) For LAM^CORE^ niches 1-5, (E) H&E images, (F) spatial mapping of cell types identified, and (G) the distributions of the cell types identified. (H) The distance of the most abundant cell types from LAM^CORE1^ in all the niches.

Niche analysis from both Xenium and Visium HD revealed similar spatially resolved expression patterns of the two LAM^CORE^ subtypes. LAM^CORE1^ cells were the primary cell type in the LAM niches, tightly clustered together at the center of the niche and surrounded by LAM^CORE2^ and lung resident cell types. This distribution suggests that within LAM lesions, LAM^CORE2^ cells at the periphery may play an important role in facilitating a favorable lesion microenvironment for the LAM^CORE1^ center through coordinated interactions with surrounding lung resident cells. These findings reinforce the idea that LAM lesions are spatially organized into heterogeneous niches, with LAM^CORE1^ and LAM^CORE2^ cells occupying distinct spatial and functional zones.

### Spatial resolution of LAM-associated fibroblast

Previously, LAFs have been reported as a population of activated fibroblasts in LAM nodules that positively express ACTA2 and CAF markers but do not express PMEL [18, 20, 21]. These cells are known for their significant heterogeneity and plasticity [14, 44]; we reasoned that the integration of single cell and spatial profiling of LAFs within their native environment would enable us to address the heterogeneity and reveal their dynamic regulation, which are fundamentally important to understanding LAM lesion formation and progression. Leveraging Visium and Visium HD with the LAM Cell Atlas and LungMAP CellRef as disease and normal lung cell references, respectively, we mapped LAM AF1 cells to distinct spatial locations, including 1) AF1 cells clustering together with LAM^CORE1^ cells but negative for PMEL and uterine specific TF (*HOXA11, HOXD11, HOXA10, EMX2*) expression (termed LAF-seed); 2) AF1 cells located in the predicted niche region surrounding the LAM^CORE1^ cells (termed LAF-niche), and 3) other scattered AF1 cells likely representing alveolar resident fibroblasts (**Fig. 8A**). As shown in **Fig. 8A-C**, LAF-seed and LAF-niche exhibit expression profiles that are clearly distinct from LAM^CORE1^ and LAM-AF1. We identified LAF signature markers and enriched functions in each region, highlighting differences compared to resident AF1 cells not adjacent to LAM niches (**Supplementary** Fig. 12). **Fig. 8D** showed the cellular composition and the expression of several LAF signature markers in a representative LAM niche.

**Figure 8:**
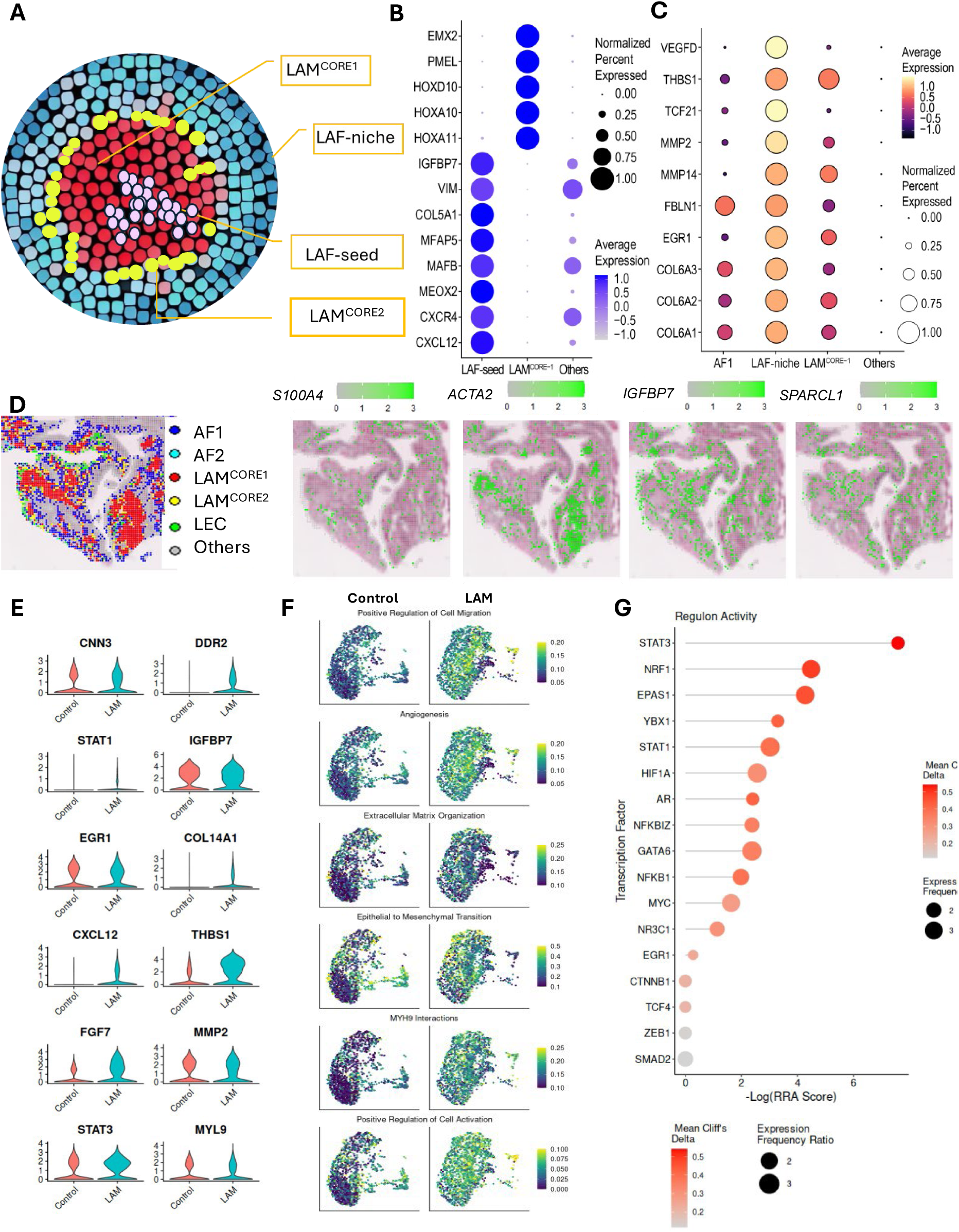
Identification and Molecular Characterization of LAFs Using Spatial and Single-Cell Transcriptomics. (A) Schematic representation of the distribution of LAM-associated fibroblasts (LAFs) and LAM^CORE^ cells within LAM niches. (B) Dot plot showing the specificity of LAF-seed marker genes compared to LAM^CORE1^ and other cell types, supporting the identity of LAF-seed cells as a distinct subpopulation. (C) Dot plot showing the expression specificity of LAF-niche marker genes, which shows considerable overlap indicating the transformed nature of the LAF-niche. (D) Left: Spatial mapping of fibroblast (AF1, AF2), LAM^CORE1^, LAM^CORE2^, and lymphatic endothelial (LEC) cells in representative LAM^CORE^-enriched niches. Right: Spatial feature plots of representative LAF marker genes (*S100A4*, *ACTA2*, *IGFBP7*, *SPARCL1*) across the same region, illustrating co-localization and spatially distinct expression domains. (E) Violin plots displaying single-cell RNA-seq expression of LAF-specific marker genes defined through spatial transcriptomics. These plots confirm that spatially identified markers are also expressed in specific AF1-derived cells, validating cross-platform consistency. (F) Module score plots for gene sets representing functional pathways enriched in LAFs based on spatial data. These include terms related to fibroblast activation, extracellular matrix organization, and wound response. (G) Top transcription factor regulons activated in LAM AF1 cells compared to control AF1 cells, as determined by SCENIC analysis. These regulons reflect LAF-specific transcriptional programs that may contribute to disease pathology and fibroblast transformation in the LAM niche.

LAF-seed and LAF-niche cells shared common functions related to cell migration, ECM organization and degradation and actively engaged in pathways related to EMT, myogenesis, muscle structure development and function, indicating their shared roles in lung injury, remodeling, and repair (**Supplementary** Fig. 12). However, LAF-seed cells were more focused on dynamic processes like cell migration, activation, response to hormone and growth factors, and stress responses; these cells were also involved in regulation of cell cycle, apoptosis, TGF-β, EMT, and Wnt signaling, further underscoring the pivotal role of LAF-seed cells in initiating changes within the LAM microenvironment (**Supplementary** Fig. 12). These cells may be highly active and capable of migrating to new sites; after arrival, they may recruit and activate additional resident lung fibroblasts to adopt LAF characteristics and form new LAM niches.

LAF-niche cells, on the other hand, had a stronger emphasis on promoting myogenesis, muscle cell differentiation and proliferation, consistent with the hallmark morphological changes in the LAM lesion, i.e., uncontrolled proliferation of smooth muscle cells. Pathway enrichment in muscle contraction, collagen and ECM degradation, and Rho GTPase signaling; over-representation of TF binding sites like SRF in the promoter regions of LAF-niche signature genes; and co-expression with smooth muscle-related genes suggests specialized function of LAF-niche cells in supporting smooth muscle cell function and proliferation (**Supplementary** Fig. 12).

We further cross-validated spatially induced LAF signatures and signaling pathways using single cell data derived from LAM AF1 and control AF1, under the assumption that signatures and pathways induced in LAF-niche should also be increased in LAM AF1 (contains LAF) vs control AF1 (lacks LAF). As shown in **Fig. 9 E-G**, genes induced in LAF-niche were also induced in LAM AF1 vs control AF1. The enriched biological processes and pathways in LAF-Niche, including EMT, myogenesis, ECM and collagen production, and angiogenesis, were induced in LAM AF1 based on the expression of associated markers and functional modules (**Fig. 9E-F**). Consistent with spatial and scRNA-seq analysis, regulon analysis of LAM AF1 vs control AF1 predicted the activation of TFs regulating genes involved in key LAF-niche biological processes and pathways (**Fig. 9G**). TFs including *ZEB1, SMAD2, GATA6, CTNNB1* and *TCF4* are known to directly involve in EMT and metastatic behavior. TFs including *STAT1, STAT3, NFKB1,* and *NFKBIZ* are central to inflammatory signaling, and the activation of these factors may play a key role in immune evasion in the LAM microenvironment. TFs include *HIF1A, EPAS1, STAT3, NRF1, MYC, EGR1,* and *YBX1* are known to regulate cell survival, proliferation, and stress response, and these factors may aid in apoptosis avoidance and stress adaptation.

**Figure 9.**
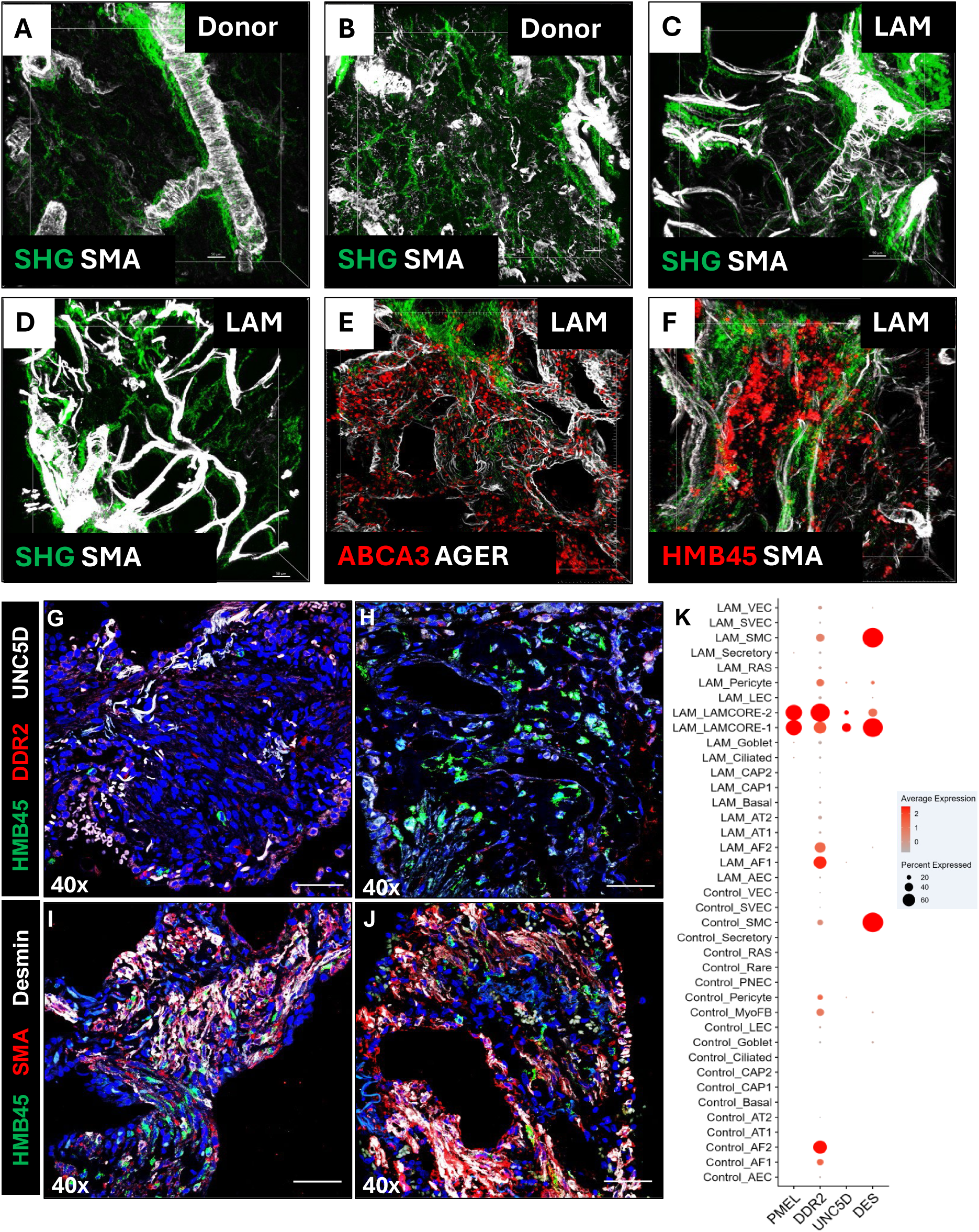
3D LAM niches visualization and LAM^CORE^ subtypes and cell heterogeneity assessment in the LAM niches. (A-B) Second harmonic generation (SHG) imaging and immunofluorescence staining for α-smooth muscle actin (α-SMA) in control lungs. (C-D) Second harmonic generation (SHG) imaging and immunofluorescence staining for α-smooth muscle actin (α-SMA) in LAM lungs. (E) ABCA3 (AT2) and AGER (AT1) staining in the LAM lungs. (F) HMB45 (PMEL) and α-SMA (ACTA2) staining in the LAM lungs. (G-H) Thin-section immunofluorescence staining of UNC5D+ (white), DDR2+ (red) and HMB45+ (green) in LAM lungs. (I-J) Triple labeling with desmin (white), α-SMA (red), and HMB45 (green) in the LAM niches. (K) Dot plots depict the PMEL, DDR2, UNC5D and DES expression across all cells in LAM and control lungs based on scRNA-seq assessment.

Overall, our approach reveals the spatial heterogeneity of LAFs and suggests a distinct biological profile associated with induced gene expression in LAFs compared to regular alveolar fibroblasts, providing new insights into their role in the disease.

### Characterize AT2 like cells in the LAM Lesion

AT2 cells represent another important niche component that is abundantly present in each of the predicted LAM niches. To better understand the alterations of AT2 cells in LAM, we identified differentially expressed genes via pseudobulk RNA-seq analysis of AT2 cells from LAM and control lungs (**Supplementary** Fig. 13A). Functional enrichment analysis of genes enriched in LAM AT2 cells revealed a significant upregulation of cytokine and TGF-β production, enhanced cell migration, cellular senescence, and DNA damage response, driven by pathways such as STAT, NF-κB, and p53 signaling (**Supplementary** Fig. 13B). Conversely, there was a notable downregulation of cholesterol and lipid biosynthesis, and phospholipid metabolic processes, likely driven by SREBP signaling (**Supplementary** Fig. 13B).

The enhanced biological processes and pathways, including cytokine and TGF-β production, cell migration, cellular senescence, mTOR, and NF-κB signaling, as well as the downregulated processes and pathways, such as cholesterol synthesis and SREBP signaling, were validated by expanding the gene pools to include all genes with the same biological terms and comparing the module scores associated with these terms between LAM AT2 and control AT2 cells (**Supplementary** Fig. 13C). Additionally, regulon analysis comparing LAM and Control AT2 cells revealed enrichment for several TFs involved in these pathways (**Supplementary** Fig. 13D).

Overall, these epithelial changes suggest that LAM alters the normal function of AT2 cells (e.g., cholesterol and phospholipid production), leading to increased secretion of pro-inflammatory cytokines that could recruit immune cells and contribute to a tumor-promoting environment.

### Experimental validation of multiomic analysis

To investigate the cellular and matrix architecture of LAM lesions, we used second harmonic generation (SHG) imaging and immunofluorescence staining on 250 µm thick lung sections from both healthy donors and LAM patients (**Fig. 9**). SHG specifically highlights fibrillar collagen, mainly types I and III, which forms organized, triple-helical bundles that provide structural support to tissues. In healthy lungs, SHG imaging revealed collagen localized around large airways (in the adventitia), within the bronchovascular mesenchyme, and as a fine network throughout the alveolar interstitium (**Fig. 9A-B**). In contrast, LAM lungs showed dense collagen accumulation around bronchovascular structures and widespread deposition throughout the alveolar regions, indicating extensive matrix remodeling (**Fig. 9C-D**).

To complement these findings, we used immunofluorescence staining for α-smooth muscle actin (α-SMA). In control lungs, α-SMA marked smooth muscle rings around major airways and a delicate meshwork within the parenchyma (**Fig. 9A-B**). However, in LAM lungs, this pattern was replaced by elongated α-SMA-positive “cables” that spanned the alveolar space. These structures, which appear as round profiles in cross-section, have traditionally been called “nodules.” Our thick-section imaging reveals that these nodules are actually cross-sections of continuous smooth muscle cables extending through the tissue (**Fig. 9C-D)**. Staining for alveolar epithelial markers, ABCA3 (AT2 cells) and AGER (AT1 cells), showed that AT2 cells remained scattered, while AT1 cells closely wrapped around the α-SMA cables (**Fig. 9E-F)**. This suggests that although the alveolar architecture is distorted, the epithelial lining is preserved. Finally, high-resolution imaging identified HMB45+ (PMEL+) LAM^CORE^ cells aligned along both collagen and smooth muscle fibers, indicating that these mutant cells localize specifically within remodeled stromal niches.

We further dissected this niche using thin sections stained for HMB45 (PMEL), UNC5D, DDR2, desmin (DES), and α-SMA (**Fig. 9G-J**). UNC5D staining outlined circular cross-sections of the α-SMA-positive cables. UNC5D (white) co-staining with HMB45 (green) was readily detectable (**Fig. 9H**) representing a population of uterine-like, LAM^CORE1^ cells. Regions lacking HMB45 signal but positive for UNC5D and DDR2 likely suggest local recruitment of lung-resident stromal cells to form these structures. Triple labeling showed desmin (DES) and α-SMA enriched at the core of the cables, with HMB45+ cells wrapping externally. In some areas, HMB45+ cells formed dense clusters lacking smooth muscle marker expression, reflecting heterogeneity in cellular composition and stromal interaction.

These structural features align with single-cell RNA expression patterns: PMEL (HMB45) is a shared marker of both LAM^CORE1^ and LAM^CORE2^ cells; UNC5D is a selective LAM^CORE1^ cells signature; DES is enriched in LAM^CORE1^ cells and smooth muscle–like cells; and DDR2 is more abundant in LAM^CORE2^ cells and fibroblast populations (AF1/AF2) (**Fig. 9K**). Thus, PMEL+/UNC5D+ regions likely represent LAM^CORE1^ cells, while DDR2+/PMEL+ zones reflect LAM^CORE2^ cells (**Fig. 9H**). Together, the data reveals a structured and molecularly heterogeneous LAM niche shaped by mutant and non-mutant cell populations and extensive matrix remodeling, including the formation of smooth muscle cables that have historically been characterized as nodules.

Immunofluorescence staining was used to validate several newly identified LAM^CORE1^ and LAM^CORE2^ cell signature genes. Hematoxylin and Eosin (H&E) staining revealed representative LAM lung nodules (**Fig. 10A-C, G, J**). The accumulation of pan-LAM marker proteins PMEL and ACTA2 was prominent in LAM lesion cells (**Fig. 10D-E**). Importantly, selective markers of LAM^CORE1^, including HOXA11 and MYH11, were abundantly present in the nucleus and cytoplasm of cells in LAM lung nodules, respectively (**Fig. 10F**). Notably, PMEL and MYH11 largely accumulated in HOXA11+ cells, which form the classical LAM nodule-like structure, indicating LAM^CORE1^ identity (**Fig. 10A-F)**. On the other hand, IGFBP7 has relatively low cell selectivity, with abundant expression in smooth muscle, endothelial and multiple other cell types. It has been reported as a secreted marker of LAF and CAF [18, 21, 45]. Immunofluorescence staining of IGFBP7 protein did not overlap with HOXA11+ cells (**Fig. 10I, N, O)** but was evident in some PMEL+ cells (**Fig. 10K, L, N, O)**, indicative of LAM^CORE2^. Our single cell analysis showed low LAM^CORE2^ selectivity for IGFBP7 alone but high selectivity when used in conjunction with PMEL (**Fig. 10P, Q, R**). Consistent with observations from spatial transcriptomics, LAM^CORE2^ cells were more scattered on the periphery of LAM niches. Taken together, immunofluorescence analyses showed distinct populations of LAM^CORE^ subtypes expressing the respective genes predicted by integrative single cell and spatial transcriptomic analyses.

**Figure 10.**
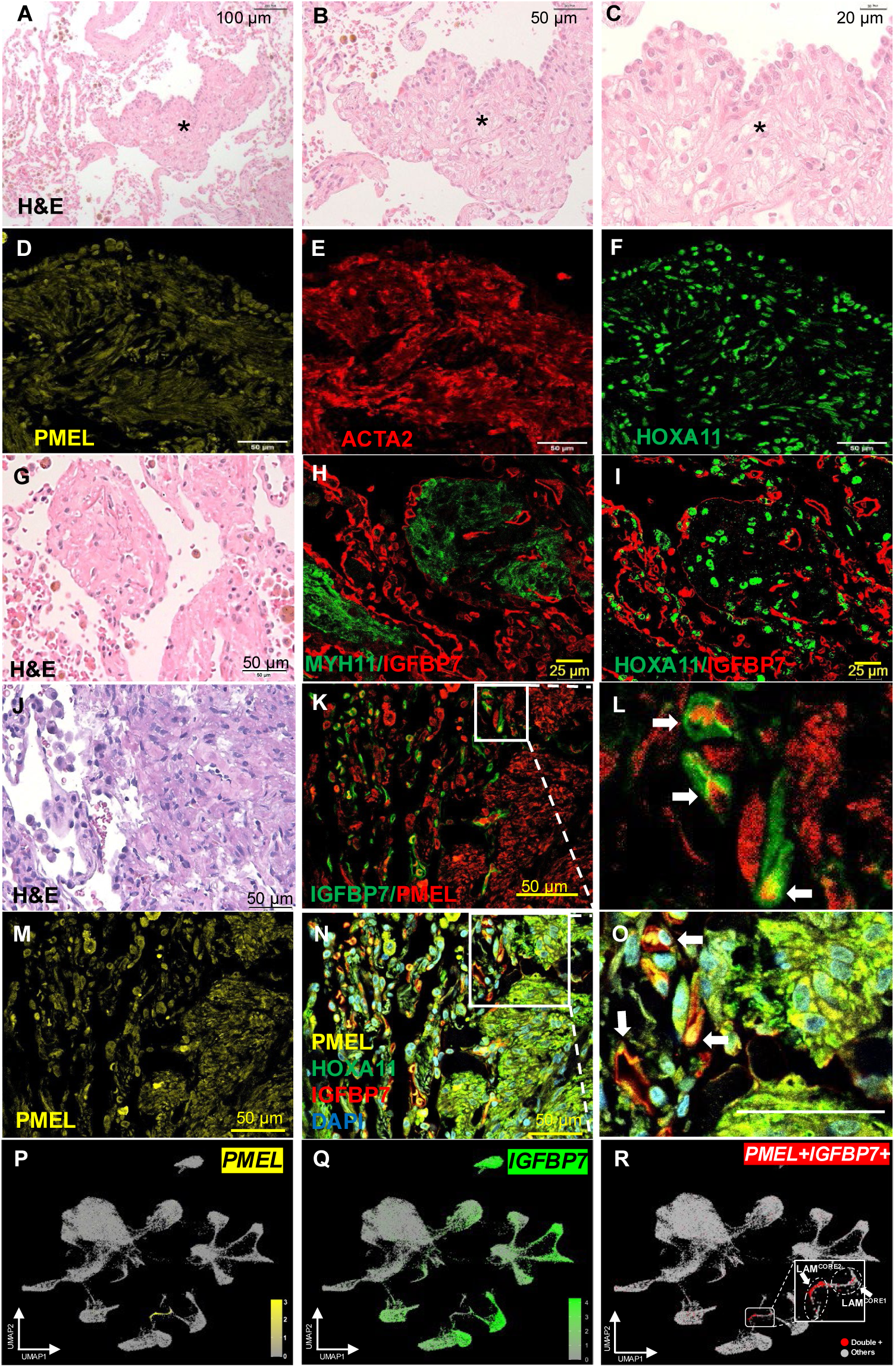
Immunohistochemistry identifies expression of genes specific to distinct cell types in LAM lung tissues. (A-C) Progressively high magnification views of a hematoxylin and Eosin (H&E) stained LAM sample. Asterisks designate LAM lesions. (D-F) LAM lung tissues were immunostained with PMEL, ACTA2, and HOXA11 and imaged by confocal immunofluorescence microscopy. (G, J) H&E-stained LAM samples (H, I, K, L) Dual staining of LAM lung tissues with (H) MYH11 and IGFBP7, (I) HOXA11 and IGFBP7, and (K, L) IGFBP7 and PMEL. (M) Staining with PMEL in LAM lung tissue. (N, O) Co-staining of LAM tissue with PMEL, HOXA11, IGFBP7, and DAPI. White arrows indicate positive staining. (P-R). Feature maps depict the PMEL+, IGFBP7+ and PMEL+IGFBP7+ double positive cells in LAM lungs based on scRNA-seq assessment.

Consistent with the immunofluorescence confocal imaging, LAM^CORE1^ cells were readily detectable by co-expression of PCP4 and PMEL or TAGLN and PMEL as determined by RNAscope in situ hybridization (**Supplementary** Fig. 14). PCP4, enriched in LAM^CORE1^, and FAP, enriched in LAM^CORE2^, were expressed in distinct cells. PCP4+ cells were more abundant, while few FAP+ cells were scattered throughout the LAM lesion **(Supplementary** Fig. 14A, C). This pattern aligns with the cell proportion and expression differences observed in the spatial transcriptome analysis between LAM^CORE1^ and LAM^CORE2^. Basal expression levels were assessed in female control lungs, revealing virtually no PMEL+ cells, while TAGLN+ cells were readily detectable **(Supplementary** Fig. 14D). The relative expression of PCP4 and FAP across all cell types were visualized in violin plots (**Supplementary** Fig. 14E). These results indicate that PCP4 and FAP are selective markers for LAM^CORE1^ and LAM^CORE2^ cells, respectively. Additionally, FAP, a CAF marker, is expressed in LAM AF1 cells at lower levels but not present in control AF1 cells, further supporting the presence of LAFs (**Supplementary** Fig. 14E).

These multimodal imaging and molecular analyses collectively revealed that LAM niches are composed of molecularly distinct, spatially organized LAM^CORE1^ and LAM^CORE2^ cells, embedded within a remodeled stromal matrix. The traditional view of LAM nodules is redefined as 3D smooth muscle cables, with LAM^CORE^ cells occupying specific niches shaped by both mutant (PMEL+) and potentially recruited stromal cells.

## Discussion

Lymphangioleiomyomatosis (LAM) is a metastatic neoplasm rather than an interstitial lung disease arising from expansion of tissue resident mesenchymal cells, as evidenced by the recurrence of pulmonary LAM following lung transplantation [1, 46]. Research from our group and others supports the uterine origin of LAM cells [12, 25, 47-49]. However, sources of cells that make up the LAM lesion and the mechanisms that control tumorigenesis, metastasis, invasion, and cystic remodeling have remained elusive remodeling [2, 46, 50]. The present study employed an integrative analysis of single-cell RNA-seq/ATAC-seq and spatial transcriptomics together with experimental validation via RNAscope, immunofluorescence microscopy, and second harmonic generation microscopy to decode the LAM niche in its native environment. The significant main findings derived from this study include: 1) the identification of two distinct LAM^CORE^ subtypes with unique signatures, functions and spatial locations, subject to divergent regulatory programming; 2) the first spatial profiling and comprehensive characterization of LAM-associated fibroblasts (LAFs) within LAM lesions; and 3) a detailed characterization of the LAM niche composition and cell-cell proximity within the LAM microenvironment.

### LAM^CORE1^ and LAM^CORE2^ Subtypes: Origins and Functional Divergence

Our study identified two distinct LAM^CORE^ subtypes, LAM^CORE1^ and LAM^CORE2^, with unique transcriptomic and functional profiles. LAM^CORE1^ cells exhibit strong similarities to uterine smooth muscle cells, expressing markers such as *ACTA2*, *MYH11*, and *MYLK*, and are enriched for pathways related to muscle contraction and uterine-specific transcriptional programs (e.g., HOX/PBX networks) [12]. These findings support the hypothesis that LAM^CORE1^ cells originate from the uterus and migrate to the lungs, where they adapt to the pulmonary microenvironment. The comparatively high entropy [33] and low pseudotime scores [51] of LAM^CORE1^ cells further suggest that these cells represents an intermediate state in this migratory and differentiation process.

In contrast, LAM^CORE2^ cells display a fibroblast-like phenotype, marked by upregulation of extracellular matrix (ECM) remodeling genes (*COL1A1*, *MMP11*, *FAP*) and pathways associated with epithelial-to-mesenchymal transition (EMT), Wnt signaling, and TGF-β activation. These cells are spatially distributed in the periphery region of the LAM niches, form more close interactions with surrounding resident lung cells in the niche microenvironment, and appear to play a critical role in modifying the lung microenvironment to facilitate LAM lesion formation. The dynamic transition from LAM^CORE1^ to LAM^CORE2^ suggests a model where uterine-derived LAM^CORE1^ cells seed the lungs, while LAM^CORE2^ cells drive ECM remodeling and niche establishment through TGF-β mediated EMT-like processes. These findings reinforce the idea that LAM lesions are spatially organized into heterogeneous niches, with LAM^CORE1^ and LAM^CORE2^ cells occupying distinct spatial and functional zones. This functional divergence highlights the need for subtype-specific therapeutic strategies, such as targeting muscle-contraction pathways in LAM^CORE1^ and ECM remodeling in LAM^CORE2^ cells.

### LAM-Associated Fibroblasts (LAFs): Key Players in Niche Formation

Our spatial and single-cell analyses revealed the presence of LAFs, which share markers with cancer-associated fibroblasts (CAFs), including *IGFBP7*, *FAP*, *S100A4*, and *SPARC*. LAFs are spatially organized into two subsets: LAF-seed cells, located adjacent to LAM^CORE1^ cells, exhibit high migratory and proliferative activity, with enrichment in TGF-β, EMT, Wnt and stress signaling pathways; LAF-niche cells, surrounding LAM nodules, promote myogenesis and smooth muscle proliferation via Rho GTPase and SRF-mediated signaling, with enrichment in angiogenesis, ECM and collagen degradation (**Supplementary** Fig. 18).

LAFs contribute to LAM pathogenesis by remodeling the ECM, secreting pro-angiogenic factors, and creating a supportive niche for LAM^CORE^ cell survival. The GRN analysis based on paired snATAC-seq and scRNA-seq profiling of LAM fibroblast identified *EGR1* as a central regulator, driving expression of activated fibroblast-associated genes (*FAP, CXCL12, and IGFBP4*), collagen genes (*COL4A1*, *COL4A2*, *CO14A1*) and inflammatory mediators (*JUNB*, *FOS, FOSL1*) (**Fig. 4**). *EGR1* is a key stress-responsive transcription factor acting as both a tumor suppressor and oncogene depending on the cellular environment. It is induced by stress signals (such as ROS, hypoxia, DNA damage), and TGF-β signaling [38, 52]. The activation of EGR1 in turn regulates the expression of various genes involved in cell proliferation, apoptosis, EMT, and inflammation. It plays a critical role in the activation of fibroblasts, promoting cancer metastasis (e.g., via MMP9 or PDGFA) and enhances tumor cell survival in certain contexts, all of which are relevant to the formation of the LAM microenvironment [38, 53-56]. These findings position LAFs as critical stromal components in LAM progression and potential targets for disrupting niche stability.

### The LAM Niche: A Unique Tumor Microenvironment

Spatial transcriptomics and high-resolution imaging delineated the cellular architecture of LAM niches, which are dominated by LAM^CORE1^ cells (32–39% of niche composition) and surrounded by LAFs, lymphatic endothelial cells (LECs), alveolar type 2 (AT2) cells, immune cells, and scattered LAM^CORE2^ cells. A key and consistent finding from the present study is the identification and characterization of LAFs, which is observed across all niches from different patients’ samples and all three spatial platforms used (Visium, Xenium and Visium HD) (**Fig. 5-7**).

**LAFs**, as a key component of LAM niches, resemble CAFs seen in many malignancies, and are characterized by increased cell activation, migration, proliferation, and cytokine expression, with profound impacts on collagen and ECM [57] (**Supplementary** Fig. 12). LAM^CORE^ cells and LAFs share common signature genes and actively synthesize and secrete ECM proteins (**Fig. 1, 8**, **Supplementary** Fig. 12). The intricate interplay between LAM^CORE^ cells (with mutations in *TSC1* or *TSC2*) and the surrounding LAFs lead to a stiff, growth-permissive pathological microenvironment in LAM and destructive cyst formation. The mTOR pathway is activated in LAM^CORE^ cells as a consequence of TSC mutations, leading to the secretion of various factors (e.g., FGF7, TGF-β, CXCL12) that can activate and recruit surrounding fibroblasts [57, 58]. In turn, activated fibroblast (i.e., LAFs) release growth factors, cytokines, and chemokines that further stimulate LAM cell growth, migration, and ECM production. This crosstalk forms a pro-proliferative and pro-fibrotic loop [59]. For example, a recent proteomic profiling of late-stage LAM lung tissue revealed significant upregulation of pathways associated with fibroblast activation and ECM deposition, including collagen deposition. Picrosirius red staining and immunohistochemistry directly confirmed increased collagen deposition within LAM nodules. This indicates a robust fibrotic response [22]. The remodeled ECM acts as a scaffold and a reservoir for growth factors, trapping and presenting them to LAM^CORE^ cells to promote cell survival and resistance to sirolimus treatment [22, 58]. While ECM deposition is crucial, LAM pathogenesis also involves dysregulated ECM *degradation*. Both LAM^CORE^ cells and LAF exhibit increased expression of various matrix metalloproteinases (e.g., MMP2, MMP9, MMP14) and cysteine protease Cathepsin K (CTSK) which degrade collagen, elastin, and other ECM components [60-62]. LAM^CORE^ cells, supported by the LAF-remodeled matrix, expand and degrade adjacent healthy lung tissue, leading to the progressive breakdown of the lung parenchyma, and continuous remodeling creates the thin-walled cysts characteristic of LAM [46, 63]. The breakdown of elastin and collagen fibers in the alveolar walls is directly implicated in this destruction.

The involvement of **LECs** is another important characteristic of LAM. In the present study, LECs closely interacting with LAM^CORE^ cells and lining the LAM niches were independently observed across different patients’ samples and from all three spatial platforms (**Fig. 5-7**), consistent with the disease’s metastatic nature. The pathogenesis of LAM is increasingly understood to involve significant lymphatic activation, with LECs playing a central role in disease progression and dissemination [2, 8, 16]. Histopathological studies have consistently demonstrated that LAM lesions are traversed by slit-like lymphatic channels lined with LECs; multiplexed imaging revealed LECs (using markers such as VEGFR3 and PDPN) closely associated with HMB45 and α-SMA-positive LAM^CORE^ cells; and the secretion and serum levels of VEGFD, a potent lymphangiogenic factor, correlate with LAM disease severity and lymphatic involvement [64-67]. Collectively, these findings establish that LEC alignment in LAM niches is not incidental but supported by extensive pathological, clinical, and molecular evidence, strongly suggesting that the lymphatic system serves as a primary route for LAM cell dissemination, contributing to the systemic nature of the disease.

**Alveolar type 2 (AT2)** cells were abundantly detected in all the niches that we randomly selected (**Fig. 5-7**). These AT2 cells within LAM lesions exhibited a marked downregulation of lipid and surfactant biosynthesis pathways (e.g., *SREBP* signaling) alongside an upregulation of pro-inflammatory cytokines, TGF-β production, and cellular senescence responses (e.g., *NF-κB*, *STAT*, and *p53* signaling) (**Supplementary** Fig. 13). This shift mirrors observations of altered epithelial cells in other conditions associated with metabolic reprogramming and immune activation. For example, in lung fibrosis, COPD, and adenocarcinoma, AT2 cells undergo senescence-associated dedifferentiation and lose their characteristic mature AT2 cell features, including surfactant production, while gaining stem-like features and senescence-associated secretory phenotype (SASP) that recruit macrophages and other immune cells and foster a pro-tumorigenic niche [68-70]. Senescent lung epithelial cells are also reported as a source of tumor-promoting factors in lung cancer [71]. A recent study demonstrated the physical interactions between metastatic cancer cells and AT2 cells within the niche. Cancer cells can prime AT2 cells into a multilineage state, and these reprogrammed AT2 cells in turn enhance the tumorigenic programs of cancer cells [68].

Notably, the upregulation of STAT, NFKB and TNF signaling and cytokine production in LAM-associated AT2 cells (**Supplementary** Fig. 13) suggests active crosstalk with epithelial and immune compartments, a feature well-documented in cancer and in idiopathic pulmonary fibrosis (IPF) [70]. In IPF and lung cancer, epithelial-derived cytokines (such as CCL2, CCL5, CXCL12, CSF1) recruit macrophages and fibroblasts, driving fibrosis and immune evasion [57, 72, 73]. Similarly, our data suggests that LAM-associated AT2 cells may contribute to niche formation through crosstalk with LAFs and LECs via paracrine signaling. The induction of senescence markers (*CDKN1A, CDKN2A*) and epithelial-derived cytokines (such as *CCL2, CCL5, CXCL12, CSF1*) in these cells further parallels the SASP observed in tumor-adjacent epithelia, where senescent cells promote chronic inflammation and tissue remodeling [57, 72, 73].

### Limitations of the study

While this study provides comprehensive insights into LAM pathogenesis, several limitations should be acknowledged. First, LAM is a rare lung disease, and the single-cell omics data were compiled from multiple institutions and platforms, posing challenges for integration and batch correction despite rigorous computational approaches. The small sample size may increase the risk of patient-specific biases. To mitigate these issues, we prioritized conserved findings across modalities and validated key results with orthogonal methods (e.g., immunohistochemistry, RNAscope). Future studies with larger cohorts, uniform platforms, and expanded spatial gene panels will be essential to confirm and extend these findings. Another important limitation is the underrepresentation of immune cells in our analysis. Since some of the publicly available single-cell RNA-seq datasets we utilized were pre-filtered to exclude immune cells (CD45-), we removed immune cells from other datasets for consistency. Consequently, our single-cell and single-nucleus analyses lack a detailed assessment of immune cell composition and function in LAM lesions. Since spatial transcriptomic data relied on single-cell references that excluded immune populations, our spatial analyses may also underestimate immune cell abundance and their potential roles in LAM pathogenesis. Future studies should employ immune-specific profiling (e.g., CITE-seq) to elucidate the immune landscape of LAM and its therapeutic implications. Finally, while HMB45 immunoreactivity, targeting the PMEL protein, is a well-established marker of TSC2-mutant LAM cells [1, 3, 74], the absence of HMB45 in some regions does not definitively exclude the presence of TSC2 mutations. These HMB45-cells may represent recruited stromal elements or potentially mutant cells with altered marker expression. Further genetic or lineage-tracing studies are required to resolve their origin and role in niche formation.

### Conclusion and Future Directions

In summary, the present study advances the understanding of LAM pathogenesis by delineating the LAM^CORE^ cell origin, heterogeneity, and the LAM^CORE^ cell enriched niche microenvironment. The identification of LAM^CORE^ subtypes and LAFs provides a framework for developing precision therapies. For example, our previous study demonstrated the use of HXR9, a HOX/PBX inhibitor, to block uterine-specific survival programs in LAM^CORE^ cells [12]. EMT or TGF-β inhibitors can be used to mitigate LAM^CORE2^-driven microenvironment remodeling and EGR1 or ECM-targeting agents can be used to disrupt LAF-mediated niche support. Future work should explore the mechanistic links between uterine LAM^CORE^ cells and pulmonary metastasis, as well as the utility of spatial technologies for monitoring niche dynamics in clinical samples. By integrating these insights, we move closer to effective, targeted treatments for LAM.

## Methods

### Sample Selection

A total of 19 lung samples were obtained from 14 female LAM patients, age 50-72, including both newly generated and previously published data [12, 21, 25-27]. Of these samples, 13 were used for single-cell RNA sequencing (scRNA-seq), 7 for single-nucleus ATAC sequencing (snATAC-seq), and 3 for spatial transcriptomics. In addition, one uterine LAM sample from a female patient [25] was included for single-nucleus RNA-seq (snRNA-seq) analysis. As a control cohort, 16 age-matched female lung scRNA-seq samples (age 51–72) were selected from the previously published CellRef dataset [28]. **Table 1** lists all samples used for this study.

### Integrative analysis of single-cell transcriptomics

RNA-seq data for 38,049 genes was analyzed. Only cells with less than 40,000 UMIs, between 500 and 7,500 expressed genes, and mitochondrial transcriptional content below 20% were retained, resulting in 74,600 LAM and 64,307 cells.

Single-cell and single-nucleus RNA-seq integration and cell clustering of the LAM samples (n=13) was done using Seurat v4.2.1 RPCA integration and Louvain clustering with a resolution of 4.6. Raw counts were normalized and scaled in accordance with the standard Seurat pipeline. The data were intentionally over-clustered to ensure within cluster purity, and clusters were manually combined into cell types based on known marker genes. Indeterminate cells were removed from further analysis. Immune cells were also excluded to remain consistent with the pre-filtered CD45-UPenn samples. Identification of LAM^CORE^ subtypes resulted from re-clustering of the mesenchymal cells alone, using Harmony v1.0 for batch correction and a clustering resolution of 1.2.

Differentially expressed genes (DEGs) were identified using Seurat’s FindAllMarkers function, with the expression frequency greater than 0.1, fold change greater than 1.5, and p-value less than 0.01. Functional enrichment was performed on DEGs using ToppFun and terms with at least 5 overlapping genes and with FDR values less than 0.1 were selected. GSEA was performed with enrichplot v1.28.1.

For LAM vs control comparisons, the control samples (n=16) subsetted from the larger CellRef integrated object and the LAM integrated object were split by sample and re-integrated using the same Seurat pipeline.

### Regulon analysis of LAM^CORE^ subtype and LAM vs control comparisons

Regulon analysis involved a combination of TF-TG interactions derived from decoupler v1.5.0 [35] and pySCENIC v0.12.1 [34, 75]. All TF-TG interactions in decoupler’s collecTRI database were used, as these were curated with experimental evidence.

Due to the stochasticity in pySCENIC, TF-TG interactions identified by the program have variations between runs. Therefore, we ran pySCENIC ten times on the integrated LAM object. TF-TG interactions that appeared at least seven times were retained and combined with the decoupler database to create a superset of regulons. Using this superset, we quantified regulon activity at the single-cell level using both pySCENIC’s AUCell method and decoupler’s univariate linear model.

We performed comparisons between both LAM^CORE^ subtypes and between the subtypes and pulmonary resident cells in LAM samples to identify regulons that were uniquely or differentially active in each subtype. For each subtype, we considered only TFs with expression frequencies of at least 10%. We used Robust Rank Aggregation algorithm [36] to consider Wilcoxon rank-sum p-value, Cliff’s delta, average ULM activity values, AUCell scores, expression frequency, and expression frequency fold change between LAM^CORE^ subtypes and between the subtypes and all other cell types. These resulted in a ranking of regulons specifically and highly active in each LAM^CORE^ subtype.

We performed similar analysis on several cell types for LAM vs control comparisons to identify unique and differentially active regulons in LAM fibroblasts and AT2 cells.

### Analysis of single nucleus ATAC-seq of LAM

Cells were obtained from snATAC-seq (n=3) and multiome (n=4) LAM samples. Cells with mononucleosomal to nucleosome-free fragments ratio less than 2, transcriptional start site enrichment score greater than 1, and fragment count within peaks between 1,000 and 100,000 were retained. Cells passing QC were integrated using Seurat V4 [76], Signac [37] and Harmony [77]. Unsupervised clustering and Harmony integration results were verified through visual inspection.

Cell type annotation of cells from the multiome samples were taken directly from the results of the integrated RNA analysis. Cells from the snATAC-seq samples were annotated by transferring cell type labels from the integrated RNA object using Seurat’s label transfer protocol. Cells labeled as immune cells were excluded to be consistent with the RNA analysis. The remaining cells were re-clustered, and clusters with fewer than 500 cells were removed due to low quality. Following a final round of clustering, clusters were annotated based on the label transfer results. Cell type annotations were validated based on the estimated activity of cell selective markers and consistency with annotations of multiome cells.

After cell cluster and cell type identification, differentially accessible peaks (DAPs) for each cell type were found using the FindAllMarkers function in Seurat, with fold change greater than 1.5 and p-value less than 0.05. The closest gene to each peak was annotated using the ClosestFeatures function in Signac. Genes that were within 10k bp of DAPs were selected for functional enrichment using ToppFun [78]. Enriched motifs were identified for each cluster by using Signac’s FindMotifs function and the DAPs of the cluster. Motif plots were generated using Signac’s MotifPlot function.

To perform peak-to-gene analysis, we extracted transcriptomic and epigenetic data of multiome cells from the integrated RNA and the integrated ATAC objects, respectively, and created a new object using Signac. RNA and ATAC assays were normalized and scaled. LinkPeaks function from Signac was used to find the correlation between peaks and the expression of nearby genes. These links, along with ATAC tracks and RNA expression were visualized using the CoveragePlot function in Signac.

### Gene regulatory analysis integrating single cell RNA and ATAC data

Gene regulatory network inference was performed using PECA2 [40] on pseudobulk RNA-seq and ATAC-seq profiles for LAM fibroblasts. Pseudobulk ATAC-seq (n=1) was obtained from subsetting data to LAM fibroblast cells annotated in the previously described integrative ATAC-seq analysis. Pseudobulk RNA-seq (n=1) was obtained by pooling expression of AF1 and AF2 cells identified by the previously described RNA-seq analysis. Edges with FDR less than 0.001 were deemed significant. The resulting network was filtered such that 1) every TG was identified as an upregulated DEG in either LAM AF1 or AF2 when compared to the corresponding control cell types and 2) every TF had an expression frequency of at least 10% of the pooled AF1 and AF2 cells.

TFs predicted by PECA2 were further compared to motifs enriched in DAPs of LAM fibroblast cells and regulons predicted for AF1 and AF2 cells as previously described. SINCERA driving force analysis [41] was performed to rank TFs based on the mean rank across six centrality scores. The top 30 TFs were selected and their edges predicted by PECA2 were visualized using Cytoscape [79]. One-hop neighbors of *EGR1* were also visualized using Cytoscape.

### LAM lung samples preparation for 10x Visium and Xenium spatial transcriptomics of Ethical approval

Samples used for 10x Xenium and Visium spatial transcriptomics were from de-identified non-used lungs donated for organ transplantation via an established protocol (approved by University of Cincinnati Institutional Review Board #2016-07095).

### 10x Visium and Xenium experiment

Tissue was collected by The National Disease Research Interchange (NDRI) from 4 donors with lymphangioleiomyomatosis lungs. Samples ≤ 0.5 cm^2^ were cut from the locations outlined above, which were fixed for 24 h in 10% neutral buffered formalin and processed into wax (FFPE). H&E staining was used to determine the morphology of tissue blocks before proceeding with ST. 10x Visium and Xenium ST experiments were performed by the Advanced Genomics Core at the University of Michigan according to the manufacturer’s protocol.

### Analysis of 10x Visium spatial transcriptomics of LAM

Visium spatial transcriptomics data were generated using the 10x Genomics Visium V4 Slide – FFPE v2 platform and processed using Space Ranger v2.1.1 with the 10x built in human reference (GRCh38-2020-A) and the Visium Human Transcriptome Probe Set v2.0. Seurat v5.1.0 was used to import Space Ranger outputs with default QC criteria to include high-confidence tissue-associated barcodes and detected genes.

Gene expression counts were normalized using SCTransform for downstream analysis. We performed principal component analysis (PCA), neighbor identification, graph-based clustering, and UMAP visualization. Differential gene expression (DGE) analysis was performed using the Wilcoxon rank-sum test. Genes with adjusted p-value less than 0.1, fold change greater than 1.5, and expression in at least 10% of cells within the cluster were considered significantly enriched in a given spatial cluster. Cells were annotated based on prediction results using RCTD [42], and human LAM Cell Atlas [27] and LungMap CellRef [28] were used as reference datasets.

### Analysis of 10x Xenium spatial transcriptomics of LAM

10X Genomics Xenium was used to perform Xenium in situ expression analysis. To perform quality control and downstream analysis, Seurat v5.1.0 package was used. Cells with between 5 and 100 features and between 10 and 400 UMIs were retained. Gene expression counts were then log normalized and scaled for downstream analysis. We performed PCA, neighbor identification, graph-based clustering, and UMAP visualization. DGE analysis was performed to identify DEGs within cell clusters. Cells were annotated based on prediction results using RCTD [42] and human LAM Cell Atlas [27] and LungMap CellRef [28] as references. Known marker genes were also used to validate cell type predictions.

Spatial niche analysis was performed using Seurat’s BuildNicheAssay function, with k-neighbors set to 20 and k-niches set to 8, to identify distinct niches based on the composition of spatially adjacent cell. LAM^CORE^ niches were then selected, and dominant cell populations within the LAM niches were explored and visualized using Xenium Explorer v3.2.0.

### Analysis of 10x Visium HD spatial transcriptomics of LAM

Cell Ranger v8.0.0 (10x Genomics) was used to align reads in FASTQ format to the human probe set and reference genome (GRCh38), producing feature-barcode matrices for analysis. We used Seurat v5.1.0 to import the cellranger count outputs. We retained barcodes with fewer than 25% mitochondrial UMIs and excluded those in the top and bottom 2.5% of gene and UMI counts to account for outliers. Using Seurat, we used SCTransform to normalize and scale the data and identified variable features, performed PCA, identified neighbors, and conducted graph-based clustering (retaining 49 principal components). We then used DGE analysis to identify marker genes within these clusters.

Spot deconvolution was used to classify and label bins with cell types derived from the LAM cell reference. We ran spacexr (RCTD) using doublet mode, which restricts any given bin to at most two cell types. The minimum UMI threshold was set to 10 due to the sparsity of the matrix and heterogeneity of the tissue. The cell types were then manually verified by comparing marker gene expressions. Using the deconvolution results, we used a custom script to identify the peripheral region (up to 50 µms from the LAM^CORE^ cells). Briefly, for each LAM^CORE^ bin, we selected all other bins within 50 µm from the outermost LAM^CORE^ cells that were not classified as LAM based on individual barcodes. We also removed any LAM^CORE^ bin that had less than 25 LAM^CORE^ neighbors, to reduce isolated LAM bins in the tissue. Then, we obtained the cell type proportions in the peripheral region and compared those proportions to cell types observed in other peripheral regions.

Visium HD typically provides spatial coordinates of each bin in pixels, i.e. pixel location based on image dimensions. Therefore, to calculate the actual distance between two bins we converted the pixel distance to the micrometer scale using the bin diameter in pixels and the image conversion factor in microns per pixel. This resulted in a Euclidean distance matrix which included the center-to-center distance of each bin. We then calculated the median distance of each cell type from LAM^CORE1^ and LAM^CORE2^. Finally, the median distances of the 10 LAM^CORE^ niches for each cell type were calculated and reported with mean proportion of each cell type.

### Experimental Validation

For immunofluorescence staining, lung sections were deparaffinized, incubated with primary antibodies (1:100 in PBS+3%BSA) and secondary antibodies (1:1,000, Invitrogen, A-21202 and A10042). Images were captured with a fluorescence microscope (Olympus BX60) or Leica Stellaris 8 Confocal Microscope.

RNAscope in situ hybridization and quantification were performed according to a protocol developed by Advanced Cell Diagnostics (ACD) [80]. In situ probes were designed by ACD. Tissue was embedded in paraffin to generate 7 μm sections. Slides were baked and deparaffinized. In situ probes were added to the slides and hybridization was performed for 2 hours at 40°C, followed by several rounds of amplification.

For fluorescence detection, opal dyes were utilized to detect the localization of the transcripts. After mounting with permanent mounting media (ProLong Gold, Thermo), slides were photographed using a confocal Nikon fluorescent microscope.

## Data Availability

Published single cell RNA-seq/ATAC-seq and spatial transcriptomic used in this study can be found in the Gene Expression Omnibus under accession codes GSE122960, GSE135851, GSE139819, GSE171524, GSE190260, and GSE217108; in the European Genome-phenome Archive under accession code EGAS00001004344; at Synapse.org under accession code syn21041850, and at LungMAP.net under accession code LMEX0000004396. New datasets generated in this study will be deposited upon publication, and scripts used for data analysis will be made available: https://github.com/xu-lab.

## Notes

### Competing Interest Statement

The authors have declared no competing interest.

### Funding

- NHLBI: R01HL153045, R01 HL172914, U01 HL175383, U01HL148856
- the LAM Foundation: LAM0150C01-22, LAM0155S01-22, LAM0164E01-1177076-24, LAMF165EI-1346848-25

