## Supplementary Figures for "Decoding Lymphangioleiomyomatosis (LAM) Niche Environment via Integrative Analysis of Single Cell Multiomics and Spatial Transcriptomics"

### Legends

**S1 – LAM Integration**
(A-D) Integrated UMAPs of 13 LAM samples displaying the distribution of (A) samples, (B) sequencing technology, (C) clusters, and (D) cell types.
(E) Bar charts showing the distributions of the samples and sequencing technology within each annotated cell type.

**S2 – LAM and Control Integration**
(A-B) Integrated UMAPs of 13 LAM and 15 control samples displaying the distribution of (A) samples and (B) sequencing technology.

**S3 – Cluster identity**
(A) Dot plot of the expression of cell type specific marker genes in each cluster. Genes were selected from LungMAP CellCards and CellRef marker gene panels.

**S4 – Cell type identification**
(A) Dot plot of the expression of cell type specific marker genes in each identified cell type. Genes were selected from LungMAP CellCards and CellRef marker gene panels.

**S5 – Quality control**
(A-C) Violin plots of three quality-control metrics – (A) number of genes, (B) number of UMIs, and (C) the percentage of mitochondrial genes expressed in each cell – with their respective cut-offs labeled in red.

**S6 – LAM$^{CORE}$ subtype unique markers**
(A-C) Top uniquely expressed genes in (A) LAM$^{CORE1}$, (B) LAM$^{CORE2}$, and (C) LAM$^{CORE3}$, as evaluated using the expression frequency of genes in each cell type. Notably, while very specific and unique markers exist for both LAM$^{CORE1}$ and LAM$^{CORE2}$, none of the top "uniquely" expressed genes for LAM$^{CORE3}$ are truly unique to that subtype.

**S7 – LAM$^{CORE1}$ regulons and target genes**
(A) Regulon activity of top LAM$^{CORE1}$ regulons and the expression of their associated target genes plotted against pseudotime. Regulons show a pattern of enrichment for muscle- and uterine-related processes.

**S8 – LAM$^{CORE2}$ regulons and target genes**
(A) Regulon activity of top LAM$^{CORE2}$ regulons and the expression of their associated target genes plotted against pseudotime. Regulons show a pattern of enrichment for muscle- and uterine-related processes.

**S9 – snATAC-seq cell type identification**
(A) Heatmap of differentially accessible peaks (DAPs) by cell type.

**S10 – LAM AF peak to gene plots**
(A) Peak-to-gene linkage plots for representative LAF genes. Multiome single-cell data integrating ATAC- and RNA-seq were used to infer correlations between chromatin accessibility at distal regulatory elements and gene expression levels, identifying putative regulatory relationships.

**S11 – Spatial mapping of LAM niches using Visium HD**
(A) H&E-level view and spatial mapping of five predicted LAM niches colored by cell types predicted by RCTD deconvolution using the LCA reference. Niche cell composition analysis revealed major cells types and their proportions within the LAM niches.

**S12 – LAF functional enrichment**
(A) Selected functionally enriched terms in LAF seed cells (top) and LAF niche cells (bottom) when compared to all other LAM AF1 cells.

**S13 – LAM AT2 characterization**
(A) UMAP plot of alveolar type II (AT2) epithelial cells subclustered from single-cell RNA-seq data.
(B) Functional enrichment analysis of differentially expressed genes (DEGs) identified from pseudobulk comparisons of LAM versus control AT2 cells. Enriched pathways reflect alterations in immune signaling, epithelial stress responses, and metabolic processes in the LAM lung environment.
(C) Module score plots for select gene sets derived from functional enrichment analysis, showing enrichment of key biological processes in individual cells. Scores reflect coordinated gene expression across pathways relevant to AT2 cell function and disease-associated reprogramming.
(D) Top transcriptional regulons activated in LAM AT2 cells compared to control AT2 cells, identified using SCENIC analysis. These regulons indicate the activation of specific transcription factor-driven programs, including immune response and cell activation.

**S14 – RNA scope validation of LAMCORE subtypes**
(A–C) RNAscope images of pulmonary tissue from LAM patients stained for: (A) *PCP4* and *PMEL*, (B) *TAGLN* and *PMEL*, (C) *PCP4* and *FAP*.
(D) RNAscope staining of control lung tissue for *TAGLN* and *PMEL*.
In (A) and (B), the co-expression of *PCP4/PMEL* and *TAGLN/PMEL* both indicate the presence of LAM$^{CORE1}$ cells (arrowheads). In (C), *PCP4* and *FAP* are expressed in distinct neighboring cells. This is expected as both are selective markers of different LAM$^{CORE}$ subtypes. In (D), control lung tissue shows negligible *PMEL* expression, while *TAGLN* transcripts are readily detected (arrowheads).
(E) Violin plots showing the selectivity of PCP4 and FAP for different LAM$^{CORE}$ subtypes.

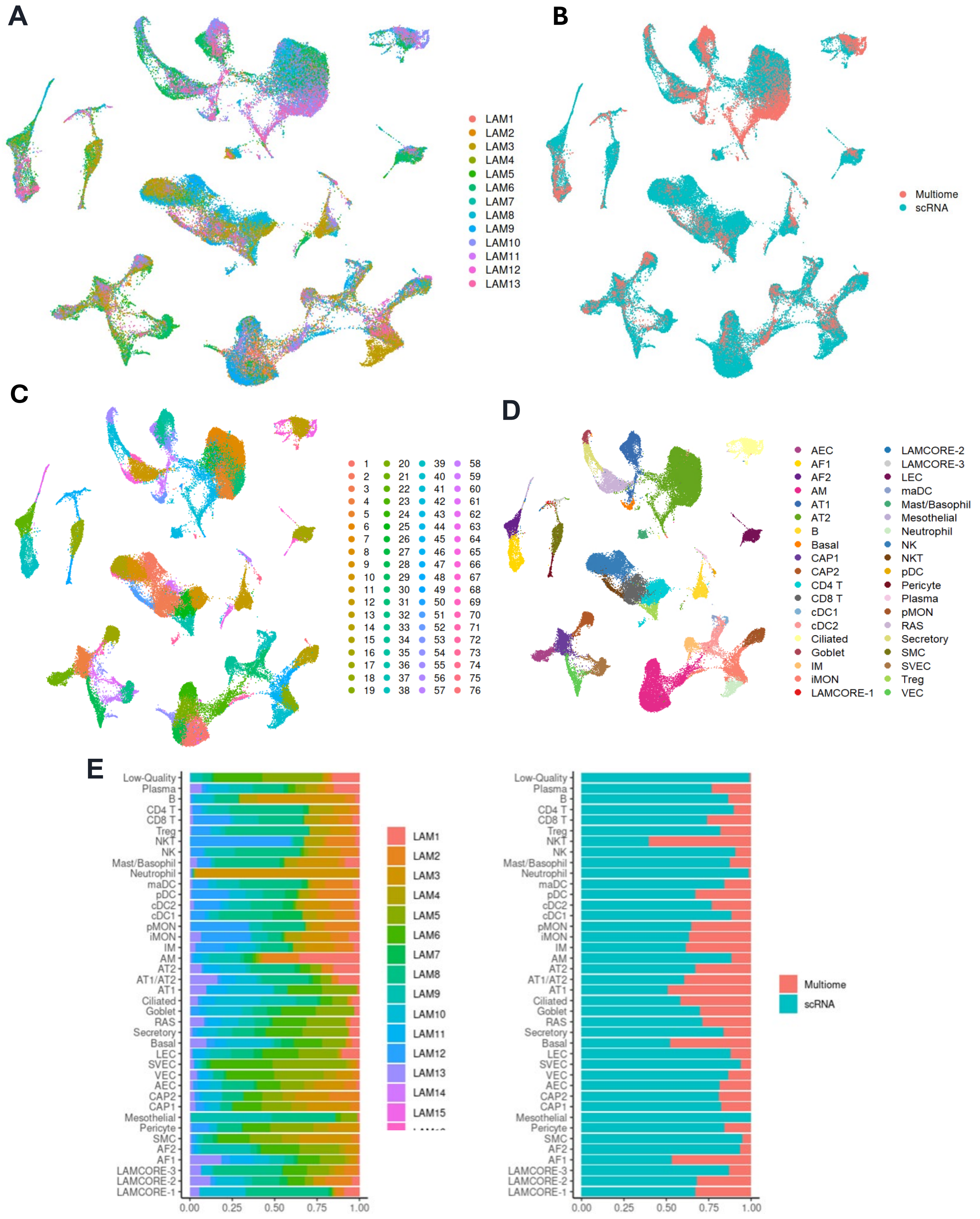


**S1 – LAM Integration**

(A-D) Integrated UMAPs of 13 LAM samples displaying the distribution of (A) samples, (B) sequencing technology, (C) clusters, and (D) cell types.

(E) Bar charts showing the distributions of the samples and sequencing technology within each annotated cell type

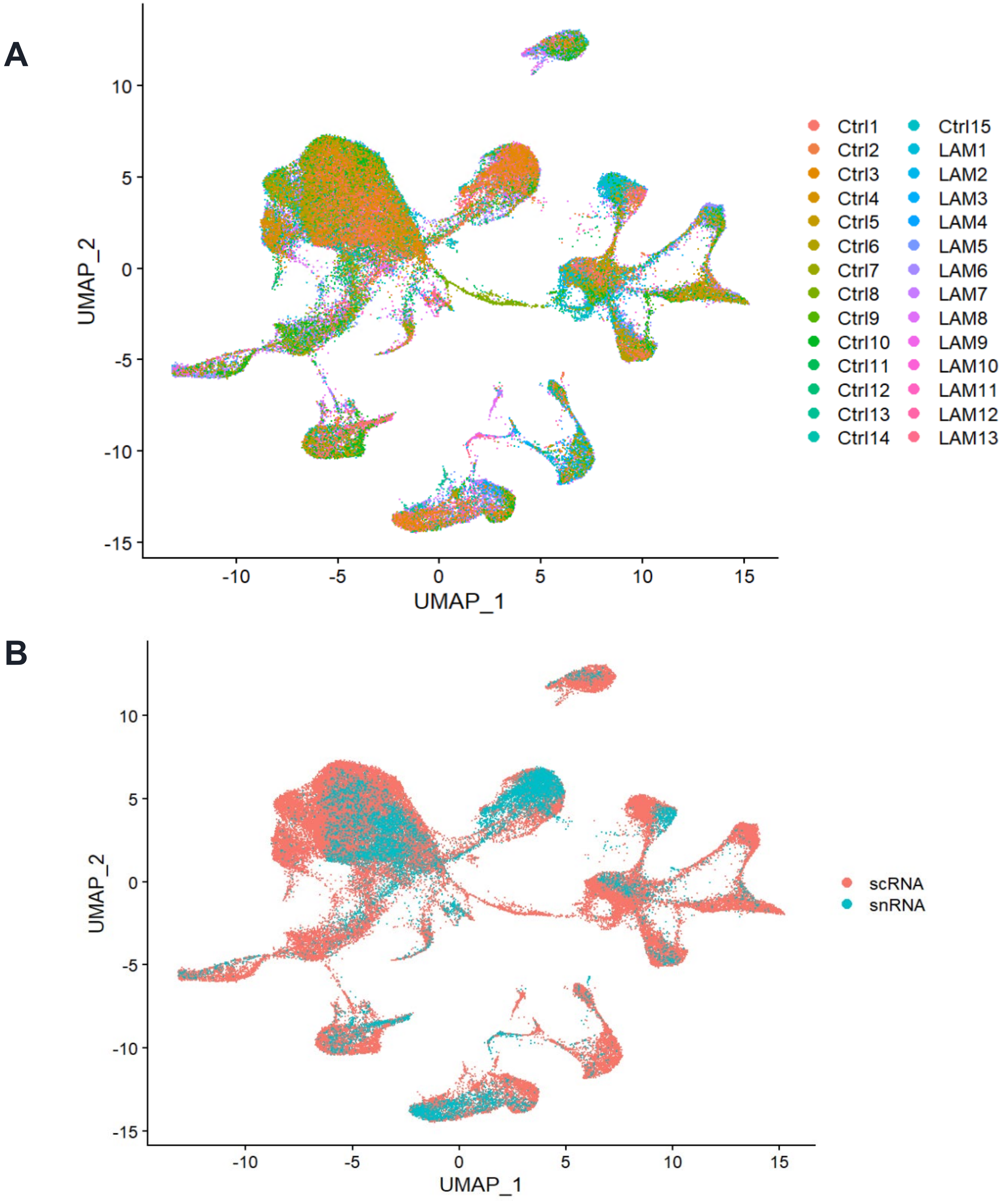


**S2 – LAM and Control Integration**
(A-B) Integrated UMAPs of 13 LAM and 15 control samples displaying the distribution of (A) samples and (B) sequencing technology.

A

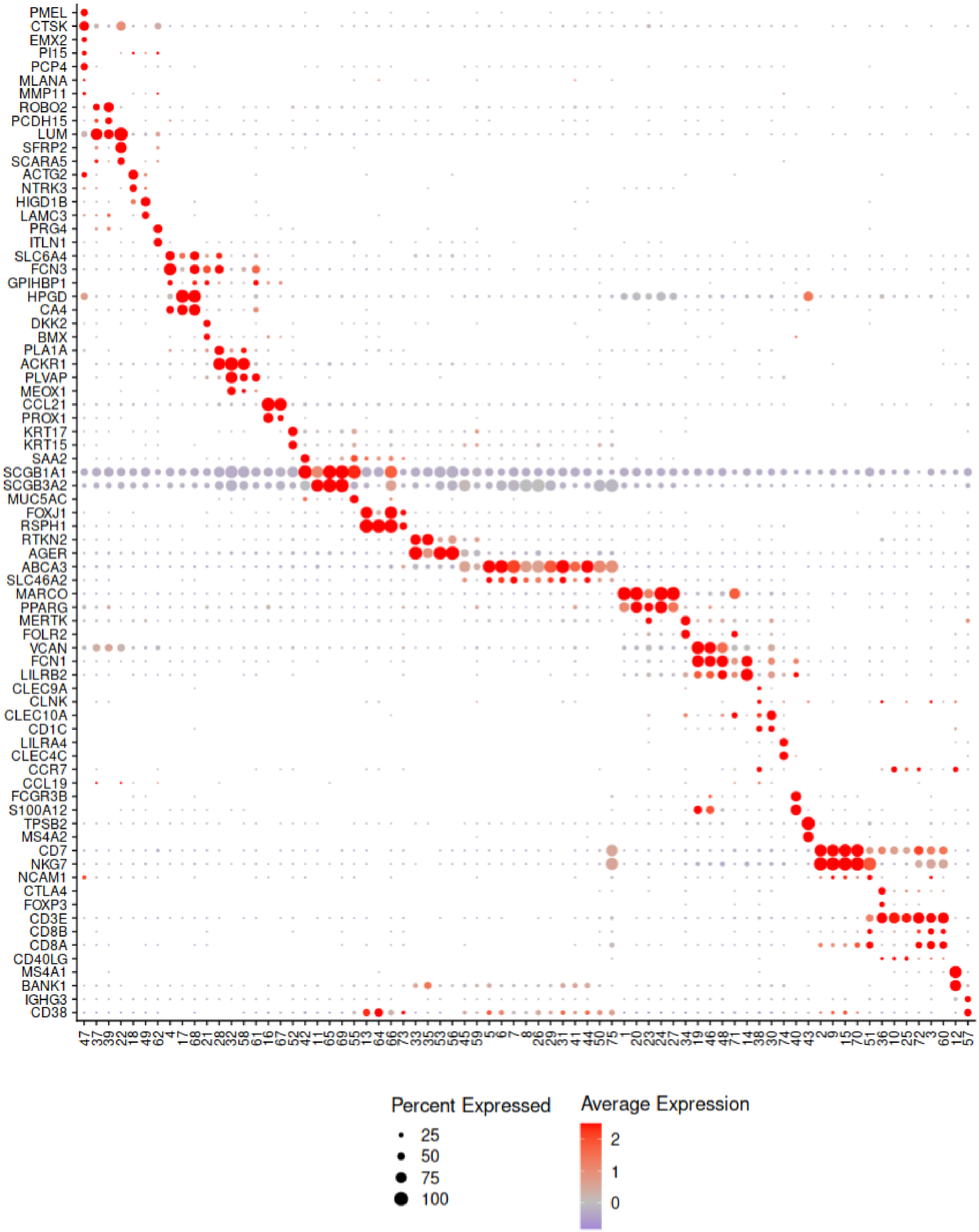


**S3 – Cluster identity**

(A) Dot plot of the expression of cell type specific marker genes in each cluster. Genes were selected from LungMAP CellCards and CellRef marker gene panels.

A

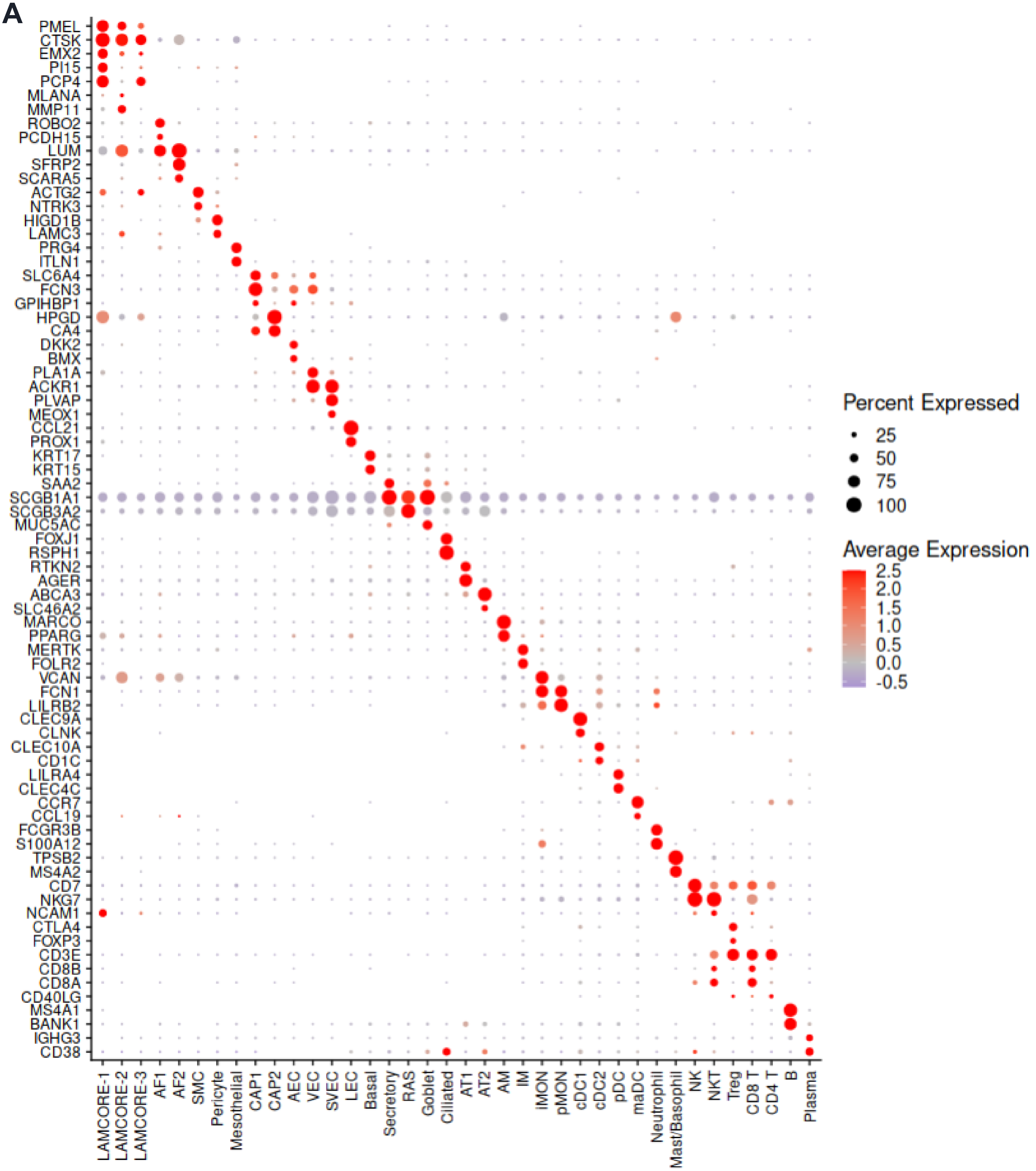


**S4 – Cell type identification**

(A) Dot plot of the expression of cell type specific marker genes in each identified cell type. Genes were selected from LungMAP CellCards and CellRef marker gene panels.

**A**

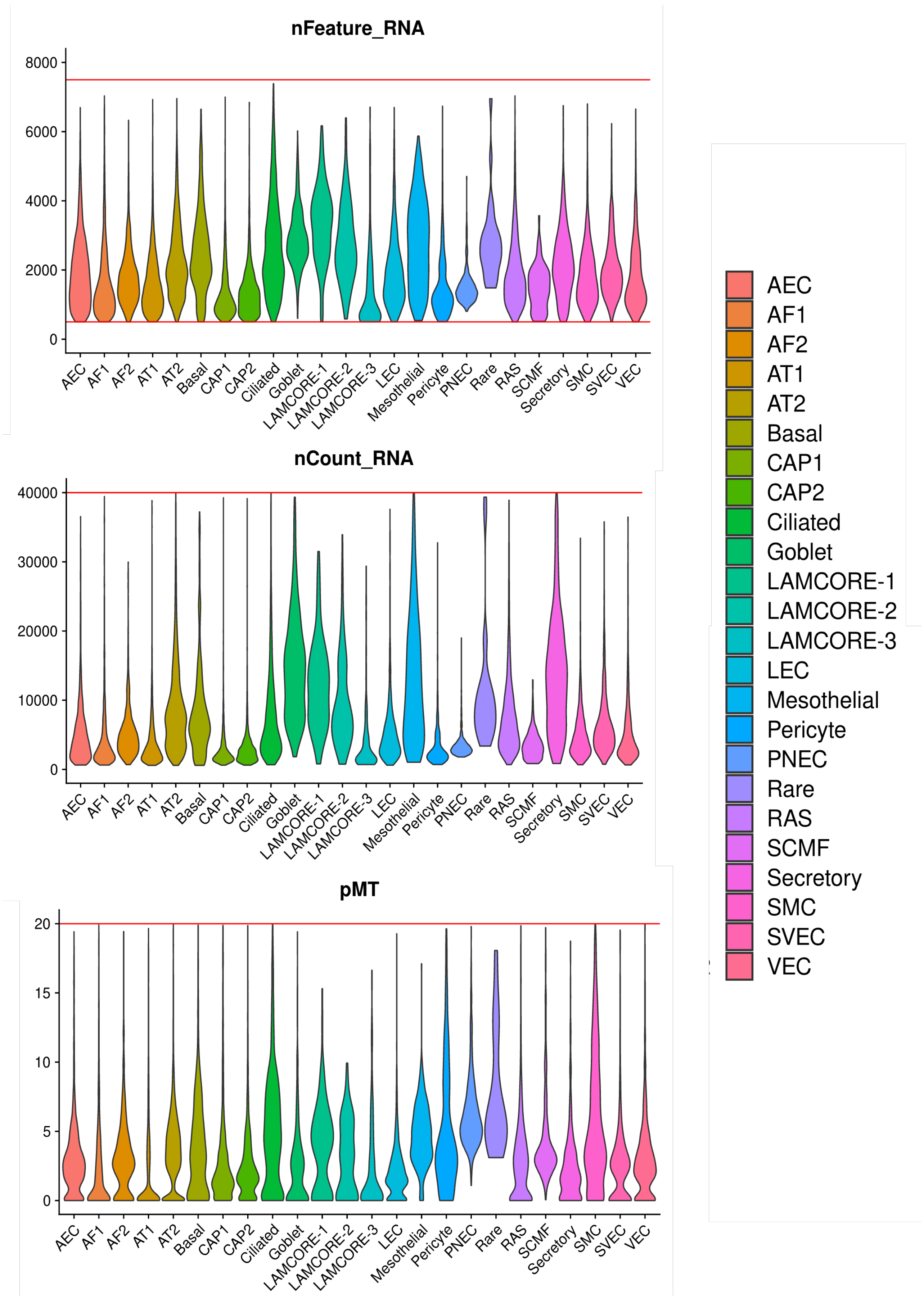


**S5 – Quality control**

(A-C) Violin plots of three quality-control metrics – (A) number of genes, (B) number of UMIs, and (C) the percentage of mitochondrial genes expressed in each cell – with their respective cut-offs labeled in red.

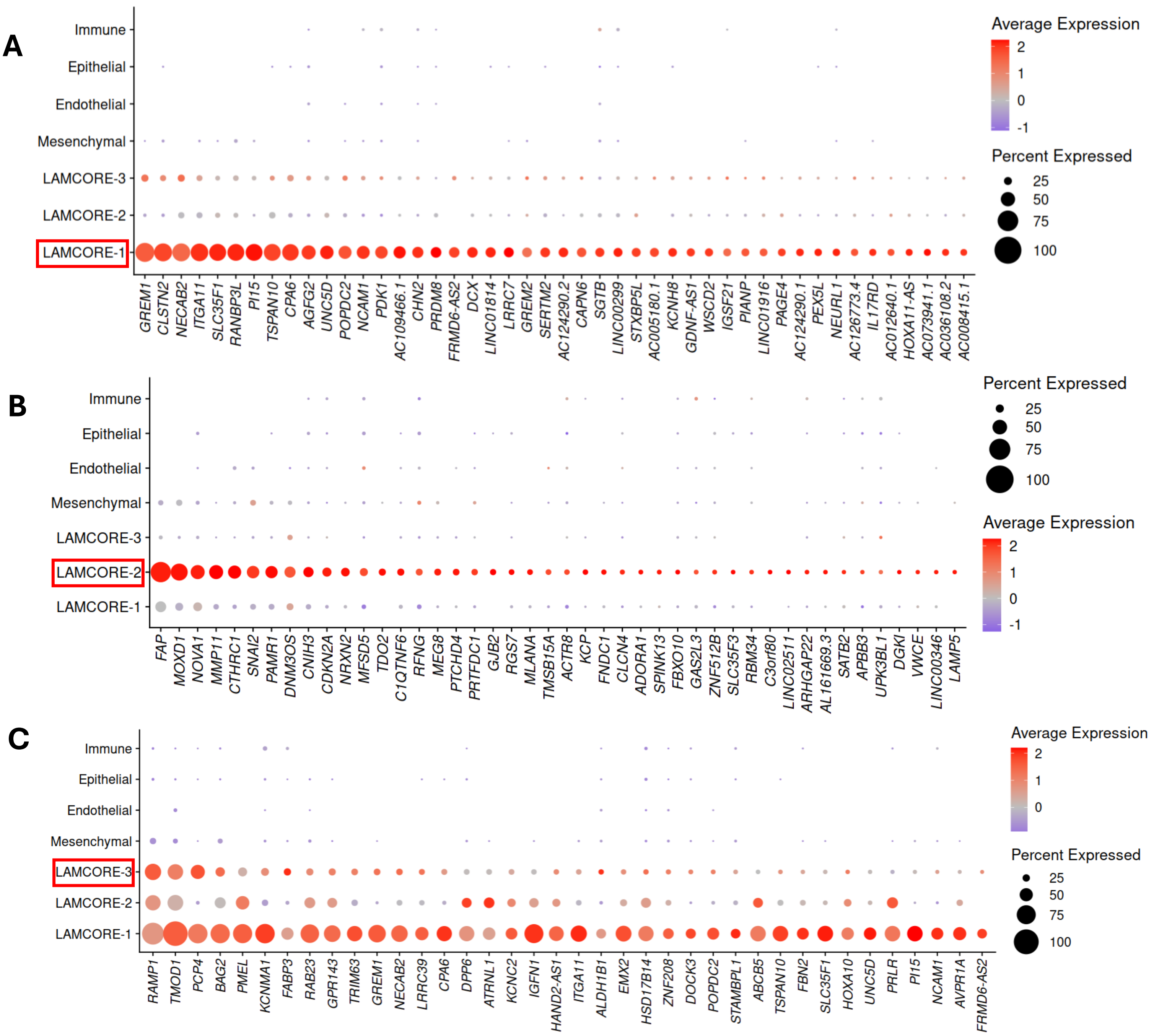


**S6 – $\text{LAM}^{\text{CORE}}$ subtype unique markers**

(A-C) Top uniquely expressed genes in (A) $\text{LAM}^{\text{CORE1}}$, (B) $\text{LAM}^{\text{CORE2}}$, and (C) $\text{LAM}^{\text{CORE3}}$, as evaluated using the expression frequency of genes in each cell type. Notably, while very specific and unique markers exist for both $\text{LAM}^{\text{CORE1}}$ and $\text{LAM}^{\text{CORE2}}$, none of the top “uniquely” expressed genes for $\text{LAM}^{\text{CORE3}}$ are truly unique to that subtype.

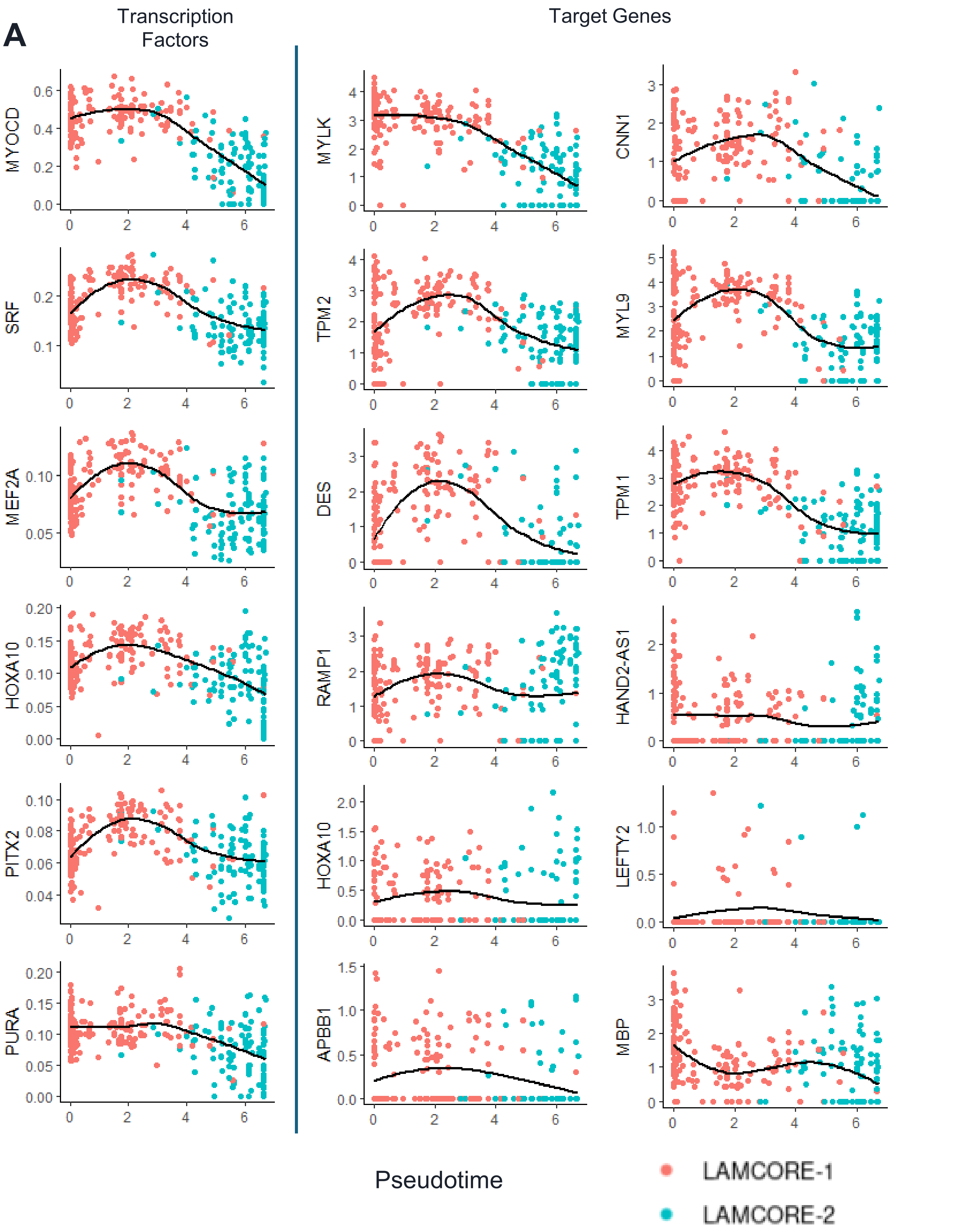


**S7 – LAM$^{CORE1}$ regulons and target genes**

(A) Regulon activity of top LAM$^{CORE1}$ regulons and the expression of their associated target genes plotted against pseudotime. Regulons show a pattern of enrichment for muscle- and uterine-related processes.

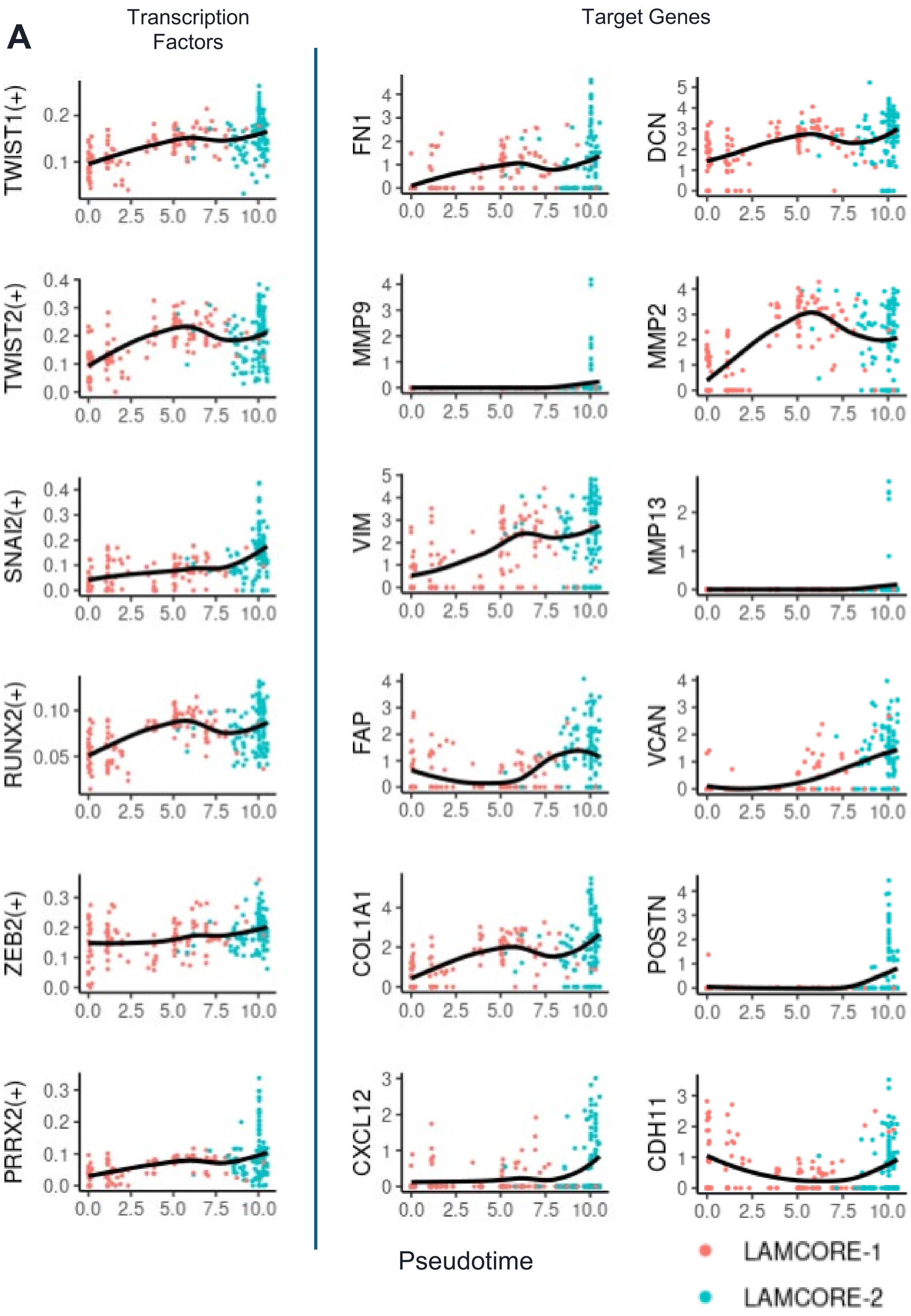


**S8 – LAM$^{CORE2}$ regulons and target genes**

(A) Regulon activity of top LAM$^{CORE2}$ regulons and the expression of their associated target genes plotted against pseudotime. Regulons show a pattern of enrichment for muscle- and uterine-related processes.

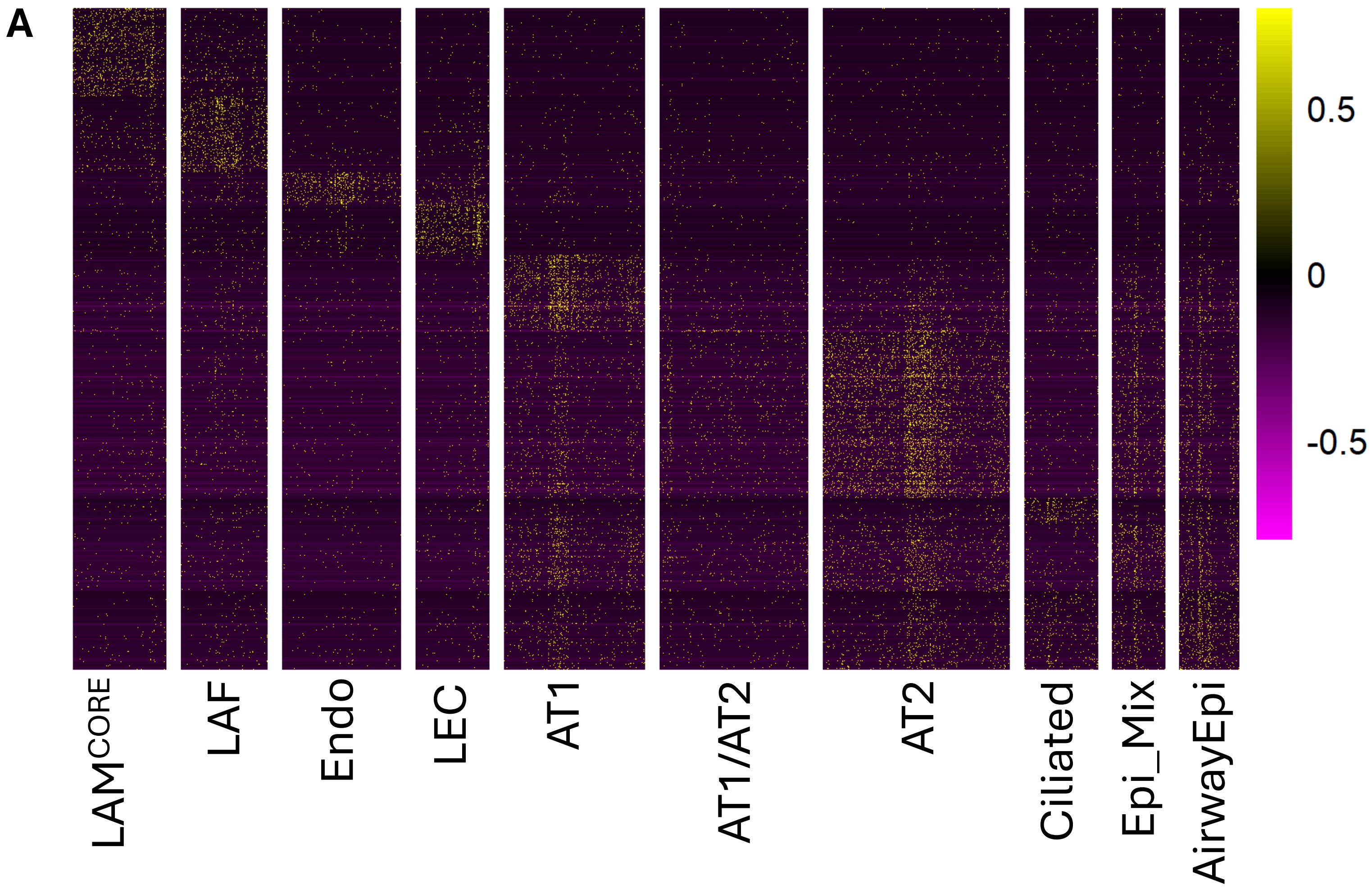


**S9 – snATAC-seq cell type identification**
(A) Heatmap of differentially accessible peaks (DAPs) by cell type.

A

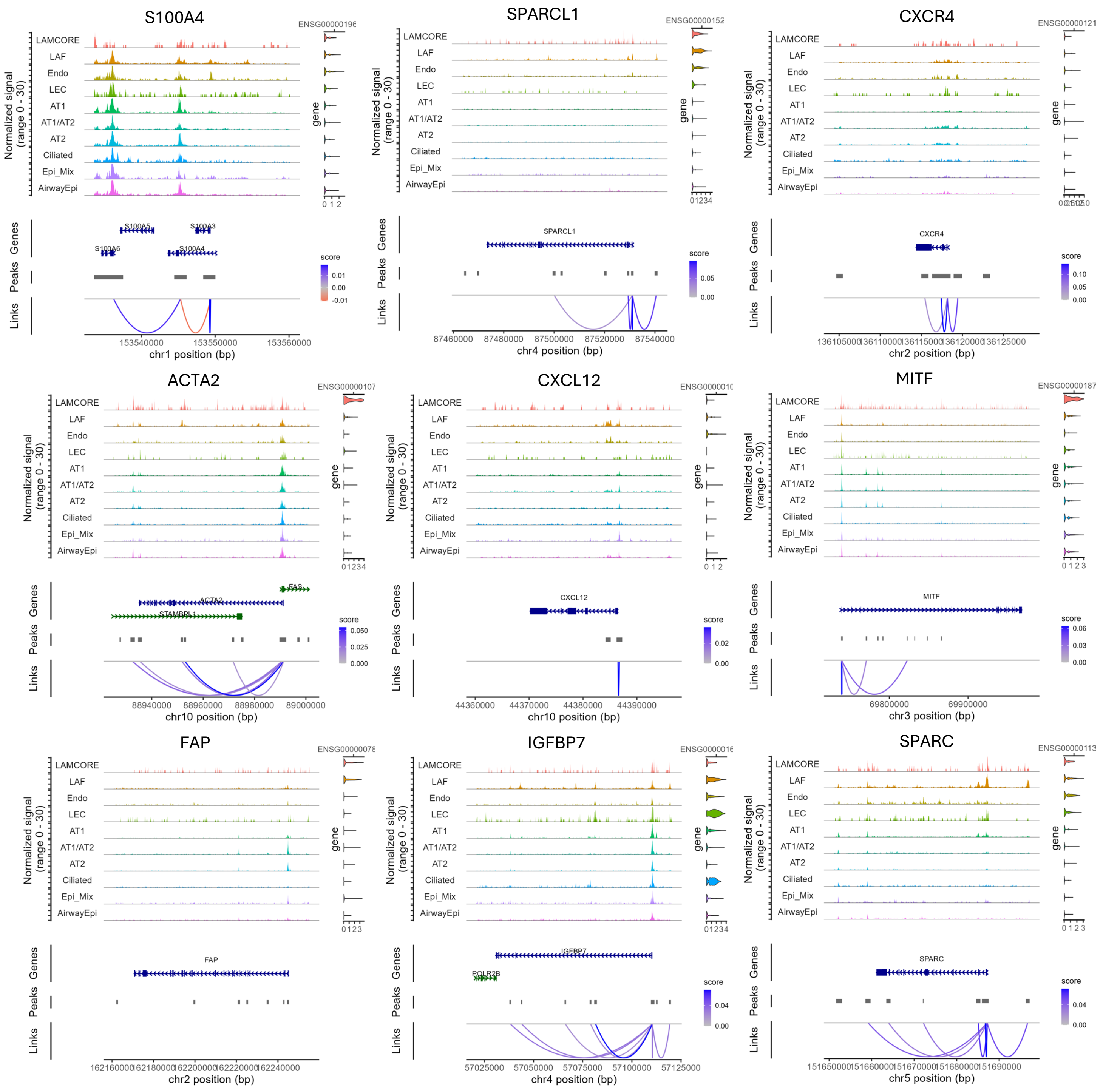


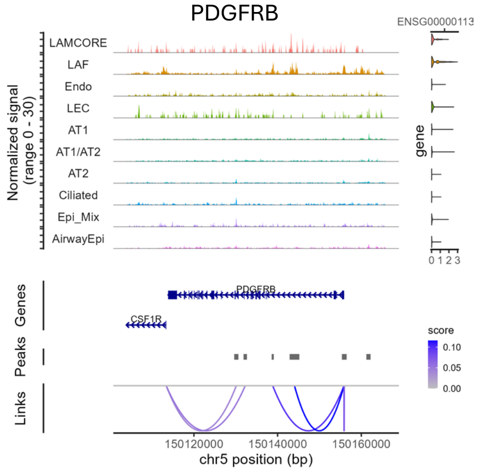


**S10 – LAM AF peak to gene plots**

(A) Peak-to-gene linkage plots for representative LAF genes. Multiome single-cell data integrating ATAC- and RNA-seq were used to infer correlations between chromatin accessibility at distal regulatory elements and gene expression levels, identifying putative regulatory relationships.

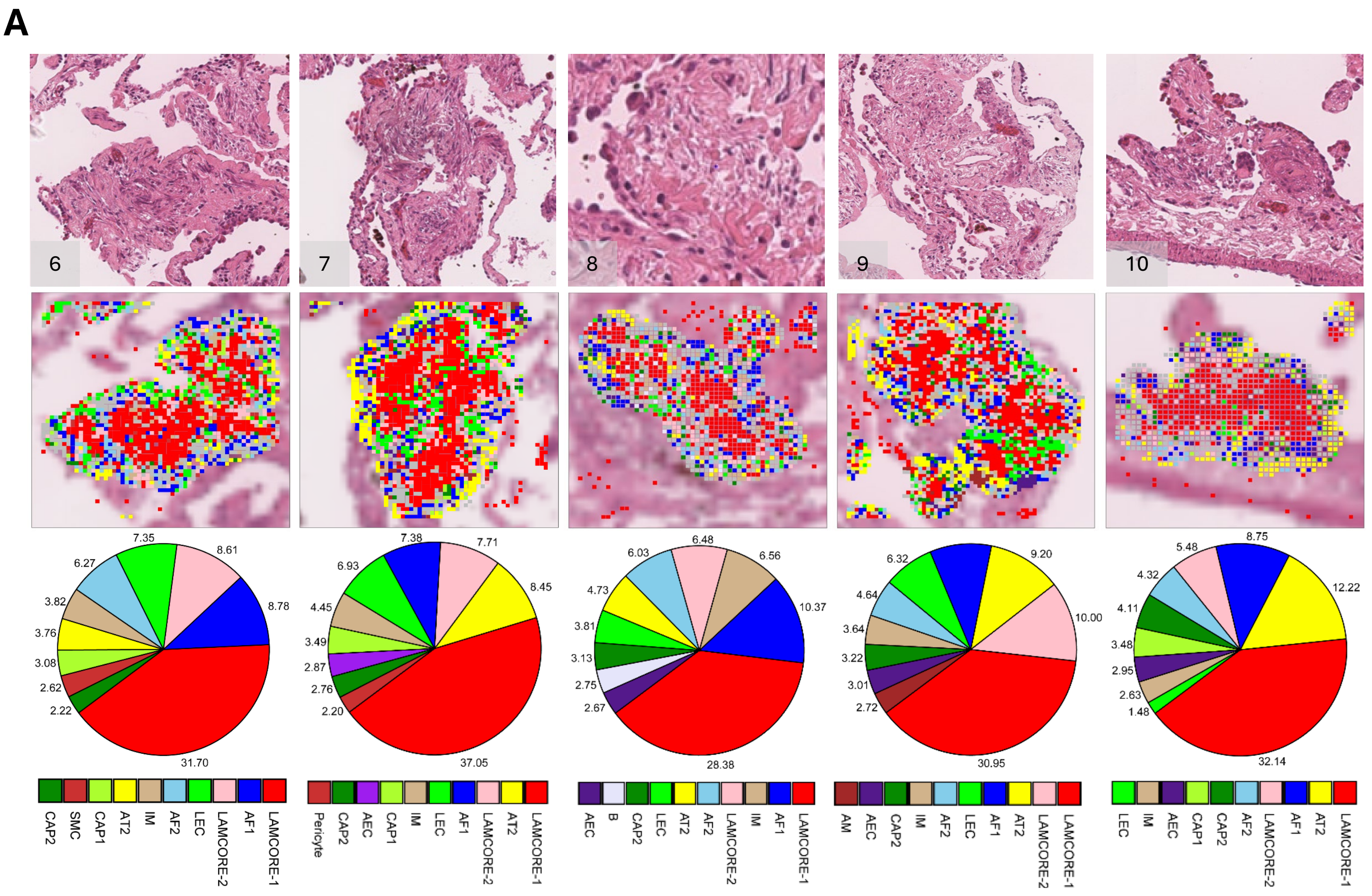


**S11 – Spatial mapping of LAM niches using Visium HD**

(A) H&E-level view and spatial mapping of five predicted LAM niches colored by cell types predicted by RCTD deconvolution using the LCA reference. Niche cell composition analysis revealed major cells types and their proportions within the LAM niches.

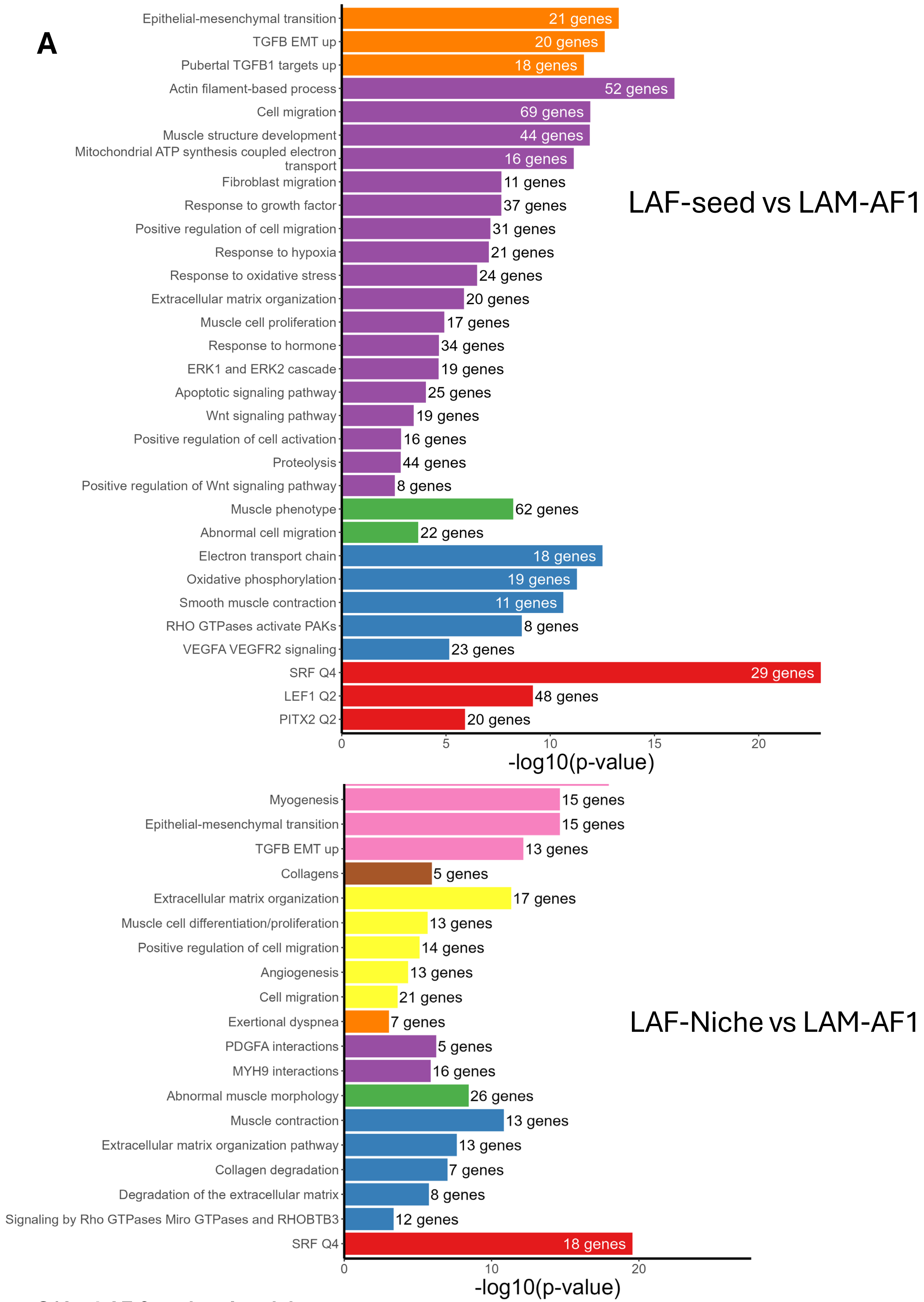


**S12 – LAF functional enrichment**

(A) Selected functionally enriched terms in LAF seed cells (top) and LAF niche cells (bottom) when compared to all other LAM AF1 cells.

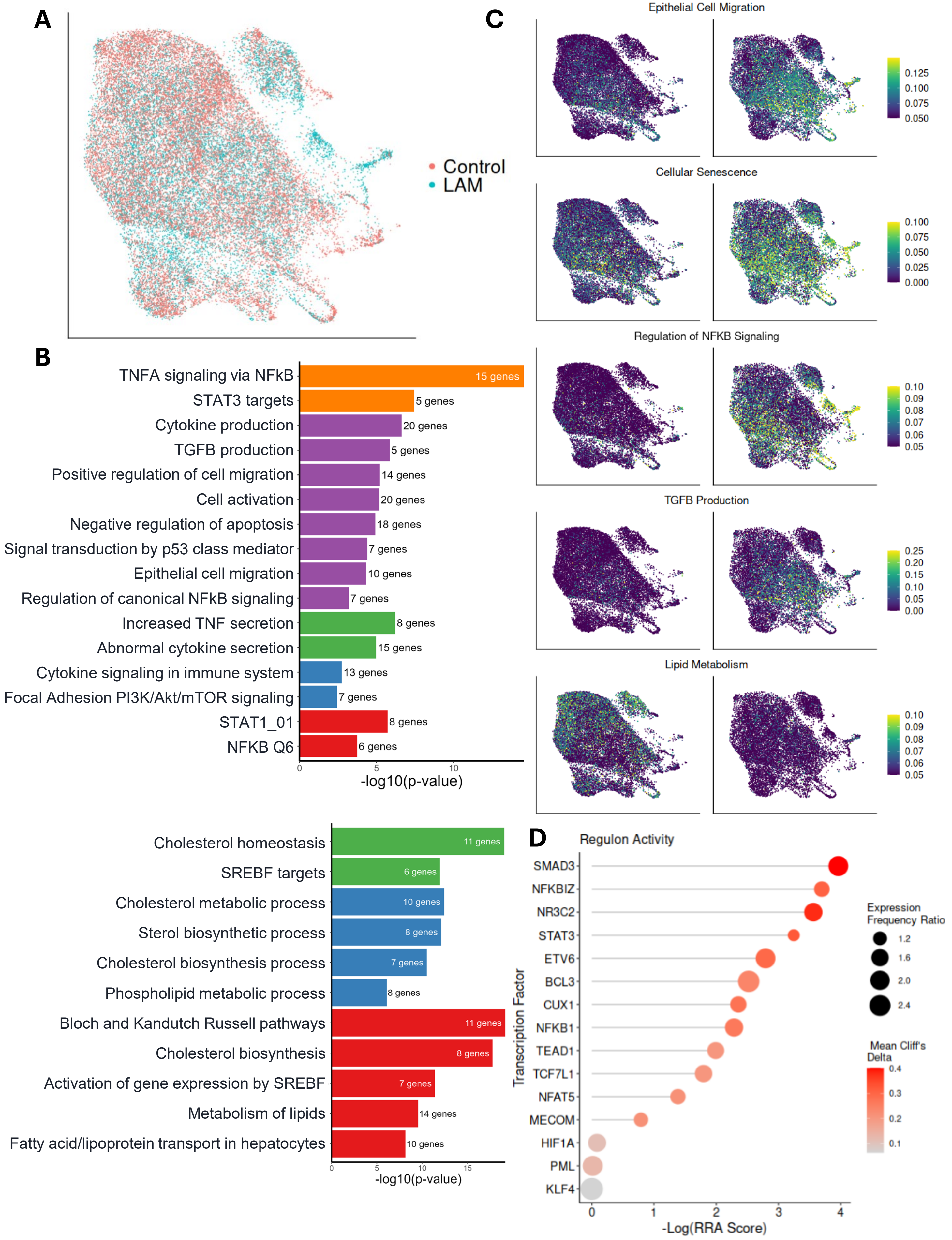
A
Control
LAM
B
TNFA signaling via NFkB
15 genes
STAT3 targets
5 genes
Cytokine production
20 genes
TGFB production
5 genes
Positive regulation of cell migration
14 genes
Cell activation
20 genes
Negative regulation of apoptosis
18 genes
Signal transduction by p53 class mediator
7 genes
Epithelial cell migration
10 genes
Regulation of canonical NFkB signaling
7 genes
Increased TNF secretion
8 genes
Abnormal cytokine secretion
15 genes
Cytokine signaling in immune system
13 genes
Focal Adhesion PI3K/Akt/mTOR signaling
7 genes
STAT1_01
8 genes
NFKB Q6
6 genes
-log10(p-value)
Cholesterol homeostasis
11 genes
SREBF targets
6 genes
Cholesterol metabolic process
10 genes
Sterol biosynthetic process
8 genes
Cholesterol biosynthesis process
7 genes
Phospholipid metabolic process
8 genes
Bloch and Kandutch Russell pathways
11 genes
Cholesterol biosynthesis
8 genes
Activation of gene expression by SREBF
7 genes
Metabolism of lipids
14 genes
Fatty acid/lipoprotein transport in hepatocytes
10 genes
-log10(p-value)
C
Epithelial Cell Migration
Cellular Senescence
Regulation of NFKB Signaling
TGFB Production
Lipid Metabolism
D
Regulon Activity
SMAD3
NFKBIZ
NR3C2
STAT3
ETV6
BCL3
CUX1
NFKB1
TEAD1
TCF7L1
NFAT5
MECOM
HIF1A
PML
KLF4
Transcription Factor
-Log(RRA Score)
Expression Frequency Ratio
Mean Cliff's Delta

**S13 – LAM AT2 characterization**

(A) UMAP plot of alveolar type II (AT2) epithelial cells subclustered from single-cell RNA-seq data.
(B) Functional enrichment analysis of upregulated (top) and downregulated (bottom) differentially expressed genes (DEGs) identified from pseudobulk comparisons of LAM versus control AT2 cells. Enriched pathways reflect alterations in immune signaling, epithelial stress responses, and metabolic processes in the LAM lung environment.
(C) Module score plots for select gene sets derived from functional enrichment analysis, showing enrichment of key biological processes in individual cells. Scores reflect coordinated gene expression across pathways relevant to AT2 cell function and disease-associated reprogramming.
(D) Top transcriptional regulons activated in LAM AT2 cells compared to control AT2 cells, identified using SCENIC analysis. These regulons indicate the activation of specific transcription factor-driven programs, including immune response and cell activation.

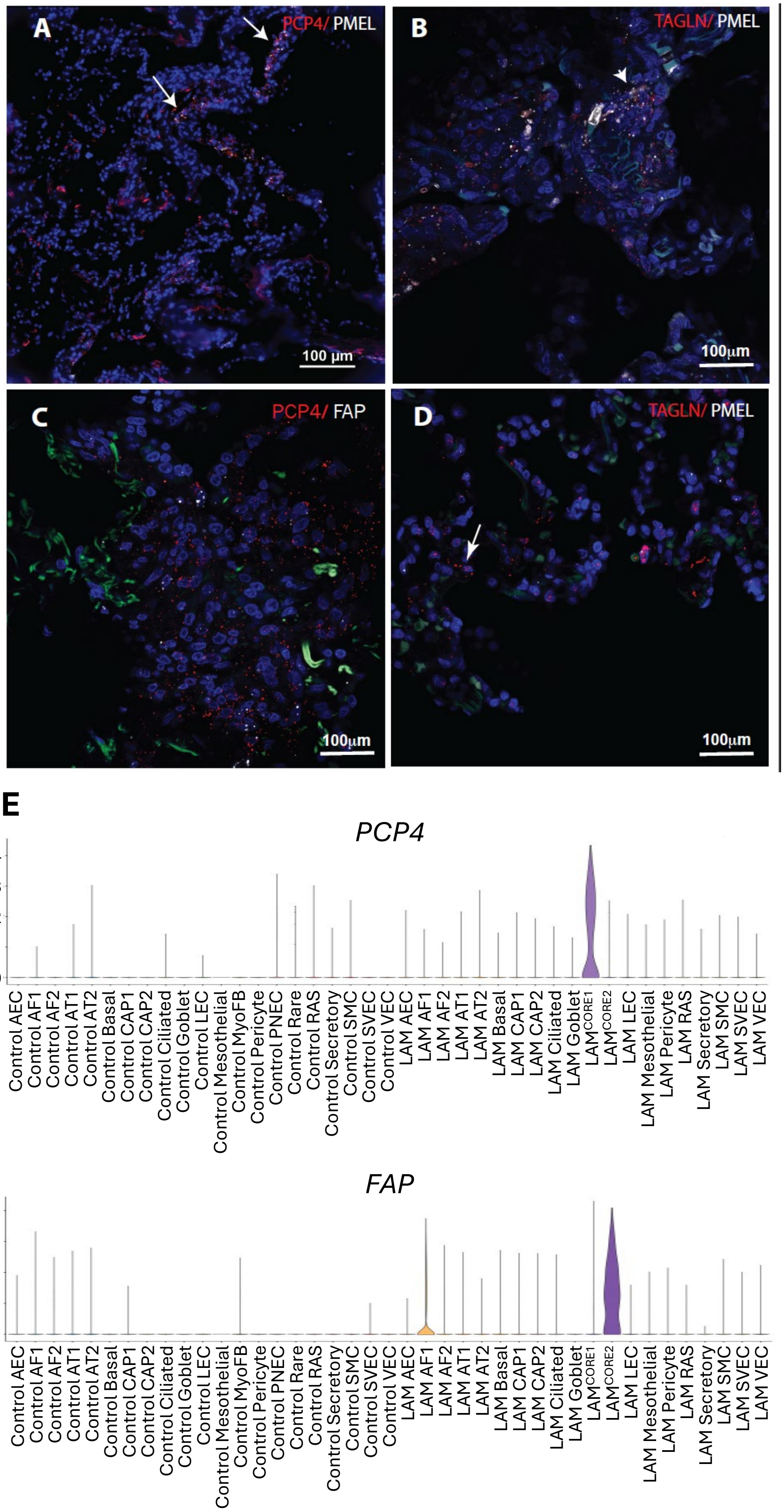


**S14 – RNA scope validation of LAMCORE subtypes**

(A–C) RNAscope images of pulmonary tissue from LAM patients stained for: (A) *PCP4* and *PMEL*, (B) *TAGLN* and *PMEL*, (C) *PCP4* and *FAP*.

(D) RNAscope staining of control lung tissue for *TAGLN* and *PMEL*. In (A) and (B), the co-expression of *PCP4/PMEL* and *TAGLN/PMEL* both indicate the presence of LAM$^{CORE1}$ cells (arrowheads). In (C), *PCP4* and *FAP* are expressed in distinct neighboring cells. This is expected as both are selective markers of different LAM$^{CORE}$ subtypes. In (D), control lung tissue shows negligible *PMEL* expression, while *TAGLN* transcripts are readily detected (arrowheads).

(E) Violin plots showing the selectivity of PCP4 and FAP for different LAM$^{CORE}$ subtypes.
